# *Candidatus* Phytoplasma-induced Retrogressive Morphogenesis in Sesame (*Sesamum indicum* L.): Tissue-Specific Metabolic and Transcriptomic Reprogramming

**DOI:** 10.1101/2025.09.29.679221

**Authors:** Saptadipa Banerjee, Gaurab Gangopadhyay

## Abstract

**Background:** *Candidatus* Phytoplasma infection is among the most destructive plant diseases, characterized by phyllody, witches broom, virescence etc. Sesame (*Sesamum indicum* L.), one of the oldest cultivated oilseed crops, valued for its nutritional and medicinal properties, is highly susceptible to phytoplasma infection, causing substantial annual yield losses. The present study aimed to investigate the metabolomic and molecular alterations in sesame plants as a consequence of phytoplasma infection in two distinct tissue types, leaves and flowers, using a multi-omics approach.

**Results:** The study revealed that phytoplasma infection significantly alters the expression of floral homeobox and meristem identity genes in an antagonistic manner, resulting in the development of leaf-like traits in floral tissues. Integrated analyses of transcriptomic and metabolomic datasets showed that control leaves, control flowers, infected leaves, and infected flowers each displayed distinct metabolomic and transcriptomic profiles. Metabolomic profiling demonstrated major changes in pathways such as porphyrin metabolism, brassinosteroid metabolism, and phenylpropanoid biosynthesis. Complementary KEGG pathway enrichment analysis of transcriptome data confirmed strong enrichment of secondary metabolite biosynthesis in both tissue types upon infection. Tissue-specific responses were evident. Floral tissues accumulated green pigments due to increased porphyrin biosynthesis and reduced degradation, while leaves showed simultaneous upregulation of both biosynthesis and breakdown pathways of porphyrins. Floral tissues exhibited stronger stress-associated responses, including upregulation of genes related to stress enzymes, phenylpropanoids, and lignification-related metabolites. In contrast, certain compounds such as lignans were specifically accumulated in leaves upon infection. These observations were further supported by biochemical, histological, and qRT-PCR assays.

**Conclusion:** This study provides the first clear evidence of tissue-specific metabolic reprogramming in sesame under Ca. Phytoplasma infection through integrated transcriptomic and metabolomic analyses. These findings improve our understanding of host-pathogen interactions and offer a basis for strategies to reduce phytoplasma-induced yield losses.

## Introduction

In angiosperms, the transition from vegetative to reproductive state is an important stage in the plant’s life and development [1, 2]. The shoot apical meristem (SAM) on perceiving certain floral induction signals, turns on the expression of flowering genes, facilitating the transition from vegetative meristem to flowering meristem [3, 4]. However, an insect-borne bacterial disease known as phyllody, caused by *Candidatus* Phytoplasma, often induces retrogressive morphogenesis, i.e., the reversion of floral tissues into vegetative structures, resulting in sterile’zombie’ plants [5]. The host range of Phytoplasmas is extensive, including plants such as periwinkle (*Catharanthus roseus*) [6, 7], tobacco (*Nicotiana tabacum* [8], maize (*Zea mays*) [9], grapevine (*Vitis vinifera*) [10], medicinal herbs such as *Atractylodes lancea* [11] and sesame (*Sesamum indicum*) [12]

*Ca.* Phytoplasma is a cell wall lacking bacteria, belonging to the class Mollicutes [13, 14] and are poorly understood as these pathogens are non-amenable to cultures [15, 16]. Phytoplasma-infected plants exhibit several symptoms like witches’ broom (excessive proliferation of axillary shoots), fasciation (flattening of stem), leaf curl, floral virescence (greening of floral organs), phyllody (leaf-like floral structures) and plant sterility. Phytoplasmas are mainly transmitted from insect vectors (leafhoppers, planthoppers, psyllids) to the plants where they thrive and multiply within the nutrient-rich phloem causing severe disfigurement and virescence of floral parts, giving a witches-broom appearances [17]. Moreover, Phytoplasma infection affects the host’s photosynthetic processes, carbohydrate metabolism, and energy metabolism, as demonstrated by physiological and molecular studies [18]. Other physiological abnormalities include chloroplast deformation and restricted transport of assimilates [19–21].

Sesame (*Sesamum indicum* L., Pedaliaceae), the present study material, is also known as the “queen of the oil seed crops”. As floral tissues get affected, infection by *Ca*. Phytoplasma is responsible for up to 80% yield loss in sesame fields [14]. Sesame phyllody disease is widely prevalent across Indian croplands, with incidence rates ranging from 10% to as high as 100% in certain regions [22]. Additionally, an early-stage infection causes seed-setting impairment, leading to considerable yield loss [10]. Recently, considering the economic importance of sesame, its global market value, and the severe yield losses caused by ‘*Ca*. Phytoplasma’ infections, several studies have focused on uncovering the underlying molecular mechanisms responsible for symptom development.

Pamei and Makandar, (2022) conducted a comparative transcriptome analysis between infected and non-infected floral buds of *S. indicum* using Suppression Subtractive Hybridization (SSH) and identified several expressed sequence tags (ESTs) associated with Phytoplasma infection. In a follow-up study, the same group performed comparative proteomic profiling of infected and healthy sesame plants. Verma *et al.*, (2022) investigated methylome alterations in healthy and Phytoplasma-infected sesame, revealing the role of DNA methylation in symptom manifestation. More recently, Verma *et al.*, (2025) and Karan *et al.*, (2025) employed transcriptomic analyses to provide insights into the molecular landscape of sesame plants infected with 16SrI-B Phytoplasmas, reporting key findings such as suppression of defense-related pathways and alterations in carbohydrate metabolism. However, no previous study on sesame has elaborated the differential impact of Phytoplasma infection on metabolomic and transcriptomic pathways across different plant tissues.

Considering the research gap, this study aims to characterize Phytoplasma-induced metabolomic changes using liquid chromatography–tandem mass spectrometry (LC–MS/MS), followed by a comparative assessment of metabolomic alterations in foliar and floral tissues upon infection. Since the degree of phenotypic alteration differs markedly between these two tissue types, we hypothesize that their defense responses will also vary significantly. Furthermore, we integrate metabolomic data with transcriptomic profiles obtained through Illumina sequencing to understand gene-to-metabolite network alterations during this unique biotic stress. The integrated data were validated through spectrophotometric assays and quantitative real-time PCR (qRT-PCR). To our knowledge, this is the first report to combine transcriptomics and metabolomics, alongside stress-related compound quantification, radical scavenging assay, antioxidant enzyme activity assays, histochemical analyses, and morphometric studies in both foliar and floral tissues, exhibiting tissue specific changes in sesame upon Phytoplasma infection. This multi-omics framework will not only provide a clearer mechanistic understanding of Phytoplasma host - interactions in sesame but also lay the groundwork for future comparative studies on molecular alterations induced by different Phytoplasma strains.

## Materials and Methods

### Site description, Experimental design and Sample collection

The Sesame plants (*Sesamum indicum* L., NBPGR germplasm collection, India, IC131989), as mentioned above, were cultivated in the Madhyamgram Experimental Farm of Bose Institute, West Bengal, India (22°41′19.68″N, 88°27′59.148″E) since 2010. Continuous surveillance of the plots showed repeated occurrence of a “witches’ broom” phenotype, which was suspected to be related to Phytoplasma infection. So, for four consecutive years (2021 to 2024), experimental plot was set, under field condition, where two identical plots were prepared each measuring 10×5 m^2^ in total isolation from other plots. Each of these plots contain ten plants with a 2-inch gap in between the neighboring plants. In one plot, the plants were sprayed with permissible dose of pesticide, as and when required, and this plot served as the control or the non-infected plants. The other plot, which was distant from the control plot, was maintained pesticide-free to promote insect vectors for a naturally occurring Phytoplasma infection. The leaf and the floral samples were collected from both the symptomatic and non-symptomatic plants, labelled, snap feezed in liquid nitrogen and stored in - 80°C until further use.

### DNA extraction and Phytoplasma confirmation

Total DNA was isolated from leaves and flowers of both symptomatic and non-symptomatic plants following the manufacturer’s protocol of DNeasy® Plant Mini Kit (Qiagen). The DNA purity was checked using nanodrop spectrophotometer (Thermo Scientific) and 1% agarose gel. For the detection and confirmation, Phytoplasma-specific universal primers P1/P7 [27] were used. The PCR condition was an initial denaturation at 94 °C for 3 min, followed by 35 cycles of 1 min at 94 °C, 1 min at 55 °C, 2 min at 72 °C, with a final extension at 72 °C for 10 min. This amplified product was used as the template for the nested PCR using the primer pair R16F2n/R16R2 [28] (Table S1), with annealing temperature 60°C and all other similar PCR parameters.

### Identification of the Phytoplasma strain and Phylogenetic analysis

To identify the Phytoplasma strain, the 1.2kb amplicon obtained as a result of nested PCR was sequenced using in-house Sanger sequencer (Applied Biosystems), following agarose gel elution using QIAquick Gel Extraction Kit (QIAGEN, Germany) according to the manufacturer’s protocol. The sequence chromatogram generated, was visualized and analysed initially with CHROMAS software (https://technelysium.com.au/wp/chromas/) to assess the quality of the data. The three replicate sequences obtained each from forward and the reverse primer were aligned using DNA Dynamo (https://www.bluetractorsoftware.com/) to generate a consensus sequence. The consensus sequence obtained was used as a query sequence to perform NCBI BLAST (https://blast.ncbi.nlm.nih.gov/Blast.cgi), selecting the organism class as “Phytoplasma”. The sequence obtained was used to perform a phylogenetic analysis.

### Evolutionary analysis by the Maximum Likelihood method

The phylogeny was inferred using the Maximum Likelihood method and Tamura-Nei (1993) model [29] of nucleotide substitutions and the tree with the highest log likelihood (-8,721.17) was generated. The tree was drawn with branch lengths (shown next to the branches) computed using the Maximum Likelihood method [30] and measured in the number of substitutions per site. The percentage of replicate trees in which the associated taxa clustered together (1,000 replicates) is shown next to the branches [31]. The initial tree for the heuristic search was selected by choosing the tree with the superior log-likelihood between a Neighbor-Joining (NJ) tree [32] and a Maximum Parsimony (MP) tree. The NJ tree was generated using a matrix of pairwise distances computed using the Tamura-Nei model [29]. The MP tree had the shortest length among 10 MP tree searches; each performed with a randomly generated starting tree. The evolutionary rate differences among sites were modeled using a discrete Gamma distribution across 5 categories (+G, parameter = 1.0817), with 6.21% of sites deemed evolutionarily invariant (+I). The analytical procedure encompassed 26 coding nucleotide sequences using 1st, 2nd, 3rd, and non-coding positions with 1,578 positions in the final dataset. Evolutionary analyses were conducted in MEGA12 [33] utilizing up to four parallel computing threads.

### Morphological and histochemical study

For this study, the symptomatic samples that showed positive amplification upon nested PCR were considered as infected samples and those without PCR amplification were treated as the non-infected samples. Stem tissues were collected from the infected and non-infected samples for morphometric and histochemical studies. For morphometric study, fresh samples were subjected to hand sectioning using a razor blade, followed by double staining using Safranin and light green. The stained sections were then visualized under light microscope and the photographs were recorded using a camera attached to the microscope (Leica).

To examine variations in secondary metabolite accumulation, free-hand cross-sections were performed and the sections were subjected to histochemical test for the detection of lignin (Phloroglucinol staining). Phloroglucinol-HCL stain was prepared by dissolving 0.3g phloroglucinol in 10ml absolute ethanol, followed by adding 5mL concentrated HCL (37%), mixed and were used for staining. Thin free-hand cross sections were submerged in Phloroglucinol-HCl stain for 5 min. The sections were rinsed with absolute ethanol to remove the excess stain, mounted in glycerol on slides with coverslips and photographs were taken.

### Sample preparation for LC-MS/MS

The leaf and flower samples of non-infected and infected sesame plants were freeze-dried using liquid nitrogen and ground into a fine powder using a sterile mortar and pestle. The powder was dissolved in 5 ml of 80% methanol (v/v), mixed thoroughly, kept overnight at 4 °C, and then centrifuged at 10,000 rpm for 10 min. The supernatant was filtered through a 0.22-µm PES filter (Millex-GP Syringe Filter Unit) for Liquid Chromatography-Mass Spectrometry (LC–MS/MS) analysis. Three biological replicates of each non-infected and infected leaf and floral tissue were used for LC–MS/MS analysis.

### Untargeted metabolomics of infected and non-infected plant samples

Liquid chromatography–tandem mass spectrometry (LC-MS/MS) analysis of metabolites was performed on a Dionex chromatography system (Thermo Fisher Scientific) coupled to an LTQ Orbitrap XL mass spectrometer (Thermo Fisher Scientific). A 5 µL aliquot of the methanolic extract was injected into a Hypersil GOLD C18 column (100 mm × 2.1 mm i.d., 1.9 µm particle size, 130 Å pore size; Thermo Fisher Scientific). The mobile phase consisted of solvent A (0.1% formic acid in water) and solvent B (0.1% formic acid in acetonitrile), delivered at a flow rate of 0.3 mL/min. Chromatographic separation was achieved over a 35 min gradient program as follows: 0 to 3 min, 75% A and 25% B; 3 to 8 min, 70% A and 30% B; 8 to 13 min, 55% A and 45% B; 13 to 23 min, 20% A and 80% B; 23 to 28 min, 20% A and 80% B; 28 to 29 min, 75% A and 25% B; and 29 to 35 min, 75% A and 25% B. The column effluent was subsequently introduced into the LTQ Orbitrap XL platform for high-resolution mass spectrometric analysis.

Consequently, obtained LC-MS/MS/MS spectra were processed and analyzed using open-source MZmine3 software (Schmid et al. 2023). Metabolites were identified using their m/z feature list obtained from MZmine. After getting entire m/z feature list, duplicate signals of adducts, isotope signals, and fragment ions were trimmed, and the final masses in the mass feature list were identified using MetaboAnalyst 5.0 server [34]. Mass intensities of each metabolite were considered for the quantitative measurement of metabolites, which are obtained from MZmine3.

For the metabolomic data obtained from LC-MS/MS, differentially expressed metabolites were determined by the log_2_ fold change and a log_2_ fold change of > 1 or < - 1 was considered as the significant. For each metabolite, log_2_ fold change was calculated using the following formula.

### Sample preparation for transcriptome analysis

Total RNA was extracted from healthy and infected samples using Spectrum plant total RNA kit (Sigma Aldrich, India) according to the manufacturer’s protocol. A total of 12 samples (3 biological replicates each of healthy and infected floral and foliar samples) were used for transcriptome sequencing. Initially, the quality and quantity were checked using NanoDrop spectrophotometer (Thermo Scientific, USA), followed by checking the RNA Integrity Number (RIN) using Agilent Tape station using High Sensitivity Screen Tape (Agilent Technologies). Following the quality control checking, mRNA was enriched using the poly-T attached beads from the total RNA, and 2×150bp paired end sequencing on the Illumina NovaSeq X Plus platform were performed by Eurofins genomic (India).

### *De novo* transcriptome analysis

Since sesame genome is not fully annotated, *de novo* transcriptome analysis was performed, after adapter trimming and removing low-quality reads using Trimmomatic v0.39 [35]. The paired end reads with phred score <25, were used for *de novo* assembly using Trinity v2.8.4 *de novo* assembler [36] with a kmer of 25 and other default parameters. The isoforms were removed from the assembled transcripts using CD-HIT-EST v4.6 and only those unigenes that have >85% coverage at 3X read depth were considered for CDS prediction using TransDecoder-v5.3.0. The filtered CDS were functionally annotated using DIAMOND [37] program (BLASTX alignment mode) that finds the against non-redundant (NR) protein database. Majority of the blast hits were against *Sesamum indicum.* Using the CDS identified, Gene ontology (GO) was performed using Blast2GO program, followed by Pathway annotation using KEGG database. The mapped result using Bowtie2, was analyzed using RSEM algorithm [38] to convert CDS abundances to FPKM. The DESEq2 package [39] was used to find the differential gene expression (DEGs) and the DEGs selected as upregulated and downregulated has fold change >1 and <-1, respectively.

## Evaluation of Secondary Metabolites

### Analysis of Total Phenolic content (TPC) and total hydrolysable tannin

To estimate the total phenolic content, 200µL of 10% (v/v) Folin-Ciocalteu reagent was added to 100 µL of methanolic extract, mixed and stored in dark for 5 min [40]. Following incubation, 500µL of 70mM sodium carbonate (Na_2_CO_3_) was added and kept in the dark for 2 hr. This reaction mixture was then used for recording the absorbance at 765nm and the total phenolic content was computed using a regression equation with gallic acid as standard. Tannins were measured by taking absorbance at 765nm and the absorbance was assessed using a UV-VIS spectrophotometer (Shimadzu) with tannic acid equivalents for preparing the standard curve.

### Estimation of Total Flavonoid content (TFC)

To determine the total flavonoid content, 300 µL of Millipore water were added to 100 µL of the 95% methanolic content, followed by adding 30 µL of sodium nitrate (5%), which were incubated for 5 min at room temperature [41]. Following the incubation, 30 µL of aluminium chloride (10%) was added and kept at room temperature for another 5 minutes. To the reaction mixture, 200 µL of NaOH (1mM) was added, mixed, and diluted by adding water to make the final volume 1mL. The reaction mixtures were allowed to stand at room temperature for 15 minutes, followed by recording the absorbance at 510 nm using UV-VIS spectrophotometer. Total flavonoid content was measured using a regression equation taking Rutin hydrate as the standard.

### Determination of chlorophyll and carotenoid

Total chlorophyll and carotenoid content was estimated by the protocol by [42]. To 100 µL of acetone extract, 900 µL of 80% acetone was added and the absorbance were recorded at 643nm, 645nm, 663 nm and 665 nm using the solvent blank. To calculate the total carotenoid content, absorbance at 436 nm was also recorded for the samples.

## Evaluation of Antioxidant activity

### Spectrophotometric Analysis of Total antioxidant assay (TAA)

Total antioxidant assay was performed using the Phospho-Molybdenum method by Prieto *et al.*, (1999). To 100 µl of the extract, 1 ml of the reagent solution (prepared by mixing 0.6M H2SO4, 28mM tri-sodium phosphate and 4mM ammonium molybdate) was added and incubated in a water bath at 90±2°C for 90min. After the reaction mixtures were cooled to normal temperature, the absorbances were recorded at 695nm against a blank mixture. The TAA concentration was measured using ascorbic acid equivalents (AAE) to prepare the standard curve.

### Spectrophotometric Analysis of Ferric-Reducing Antioxidant power (FRAP)

The FRAP was determined using the protocol of Benzie and Strain, (1996). The FRAP reagent was prepared by mixing 2,4,6-tripyridyl-s-triazine (TPTZ) (10mM (w/v), prepared in 40mM HCL; 20mM FeCl_3_ and 300mM sodium acetate buffer (pH 3.6) in a ratio 1:1:10. To 100µl of sample extract, 900µl of pre-warmed FRAP reagent was added and were incubated at 37°C for 5 min to develop a blue colour complex. The absorbances was measured at 594nm in UV-VIS spectrophotometer and total FRAP was calculated using different concentrations of Ferrous Sulphate (FeSO_4_) for the regression equation at A_594_.

### Spectrophotometric analysis of DPPH radical scavenging activity

To determine the DPPH radical scavenging activity of the extracts, 1.2ml DPPH solution (w/v, prepared in methanol) was added to 100µl of the extract, followed by a dark incubation for 15 min at room temperature [45]. The decrease in absorbance was measured at 517nm using ascorbic acid as standard. The control samples were the reagent mixture without any sample. The percentage scavenging activity was calculated using the following equation:

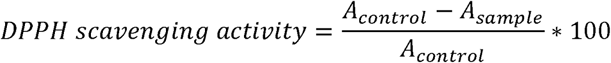

where

A_control_ = Absorbance of the control; A_sample_= Absorbance of the sample

The DPPH scavenging activity in the samples was calculated from the calibration curve using ascorbic acid as standard and are expressed as % of scavenging activity/g of tissue.

### Spectrophotometric analysis of ABTS radical scavenging activity

The ABTS radical scavenging assay was performed according to the protocol by <u>Re *et al.*, (1999)</u>. To prepare the reagent, 7mM ABTS (2,2′-azino-bis (3-ethylbenzothiazoline-6-sulfonic acid), dissolved in water) was mixed with 2.45mM Potassium persulfate (K_2_SO_4_) and was allowed to stand undisturbed in dark at room temperature for 16 h. To prepare the working solution, after 16 h of incubation, the ABTS reagent was diluted by adding ethanol to achieve an absorbance of 0.70±0.02 at 734nm, and this solution served as the control. To 100µl of the sample extract, 1 ml ABTS reagent was added, followed by dark incubation for 5min. The colour change was observed at 734nm in comparison to ethanol as blank. The percentage scavenging activity was calculated using the following formula:

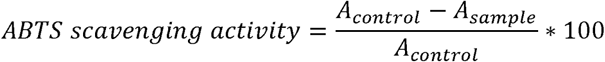

where

A_control_ = Absorbance of the control; A_sample_= Absorbance of the sample

The ABTS scavenging activity in the samples was calculated from the calibration curve using the common antioxidant, ascorbic acid, as a standard and are expressed as % of scavenging activity/g of tissue.

### Assessment of Superoxide (SO) scavenging activity

Superoxide scavenging activity was measured according to (eauchamp and Fridovich, (1971). To 100μl of the methanolic extract, 1.8ml phosphate buffer (pH 7.6), 20μl of 2.66mM riboflavin and 80μl of EDTA (12ul) was added. To this 100 µl NBT (of 1.22nM) was added, followed by light incubation (20W white light bulb) for 90 seconds at room temperature in comparison to a blank which is the non-illuminated reaction mixture. Results were calculated using a regression curve of quercetin as standard.

### Antioxidant enzyme activity assays

#### Preparation of crude enzyme extracts and estimation of protein

To prepare the crude extract, 1gm of fresh sample was finely crushed using liquid nitrogen and dissolved in 5mL of 0.1 M potassium phosphate buffer (pH 7). The mixture was centrifuged at 16000 rpm at 4°C for 20 min and the supernatant was collected for all the assays. Total protein content was estimated using the Bradford technique [48].

#### Assay of antioxidant enzymes

Guaiacol peroxidase activity was estimated following the protocol by Moon and Mitra, (2016). To 650 µL of phosphate puffer, 100 µL of crude enzyme extract was added, followed by adding 150 µL of 20mM guaiacol and 100 µL of 1 mM H_2_O_2_. The product concentration was recorded at 436 nm using UV-VIS spectrophotometer and the concentration was calculated using the molar extinction coefficient of 25500 M cm^-1^.

To determine the Ascorbate peroxidase activity, the protocol by Jebara *et al.*, (2005) was followed. For APOX activity estimation, 100 µL of the crude extract was added to the reaction mixture, which consists of 790 µL phosphate buffer, 10 µL 0.5mM ascorbic acid and 100ul of 1 mM H_2_O_2._ The concentration was measured at 290 nm and the molar extinction coefficient of 2.8 M cm^-1^ was used for enzyme activity quantification.

Polyphenol oxidase (PPO) activity was measured by using the protocol Chung and Felton, (2011). For PPO activity estimation, 10 µL of crude extract was added to 200 µL of 3mM caffeic acid. The reaction results were monitored for 5 min and was calculated using the molar extinction coefficient of 2062 M cm^-1^.

### Gene expression analysis using qRT PCR

Replica aliquots of RNA samples, used for the transcriptomics analysis, served as the templates for validation experiments. The qRT-PCR was performed using Maxima SYBR Green/ROX qPCR mix (Thermo Scientific) in an AriaMx real-time PCR (Agilent) machine after the synthesis of cDNA using QuantiTect® Reverse Transcription kit, following the manufacturer’s protocol. For a single qRT-PCR reaction mixture (25µL), 12.5 µL of 2X master mix, 9.5 µL of nuclease free water, 0.5 µL of forward and reverse primer and 2 µL template were mixed. In this study, a few essential genes from each pathway that were enriched from both metabolomics and transcriptomics data were selected and subsequently validated (Table S2). Three independent biological replicates were used and the fold Change in gene expression was calculated using the formula of 2^−ΔΔCt^ [52] using actin as a reference gene.

## Statistical analyses

All morphometric and spectrophotometric data were analyzed using Student’s t-test (control vs. infected) to determine statistical significance, with a threshold of 95% confidence (p < 0.05). Leaf (control and infected) and flower (control and infected) samples were treated as independent groups during statistical testing.

For omics datasets (metabolomics and transcriptomics), multivariate statistical approaches were applied. Principal component analysis (PCA) was performed for ordination analysis to evaluate the clustering and separation of samples based on overall metabolite and transcript profiles. For pathway impact analysis of metabolomic data (in metaboanalyst 6.0), differentially accumulated metabolites (DAMs) from leaf and flower tissues were mapped onto Arabidopsis thaliana ortholog-based KEGG pathway maps. Pathway topology analysis was then performed by integrating over-representation statistics (enrichment scores) with pathway structure-based metrics such as betweenness centrality, thereby quantifying the relative impact of each pathway and identifying the most significantly impacted metabolic routes. For transcriptomic data, enrichment analysis was carried out using ShinyGO. Differentially expressed genes (DEGs) were uploaded and queried against the Sesamum indicum gene library. The analysis combined gene set enrichment statistics with functional annotation clustering to identify significantly overrepresented KEGG pathways.

Unless otherwise specified, all statistical analyses were conducted in OriginPro 2023, which was also used for all graphical representation of results.

## Results

### Confirmation of Phytoplasma Infection using nested-PCR

The non-symptomatic (Fig. 1a) and symptomatic plants (Fig. 1b-c) were checked for Phytoplasma infection using nested PCR process. In the first round of PCR with universal Phytoplasma primers, P1/P7, a ∼1.6 kb amplicon was detected exclusively in symptomatic samples, while there was no amplification in the non-symptomatic plants (control). The subsequent nested PCR using R16F2n/R16R2 primers produced a _∼_1.2 kb product only in the symptomatic plants, thereby confirming the presence of Phytoplasma 16S rRNA and validating infection in the infested sesame plants (Fig. 1d-e).

**Fig. 1:**
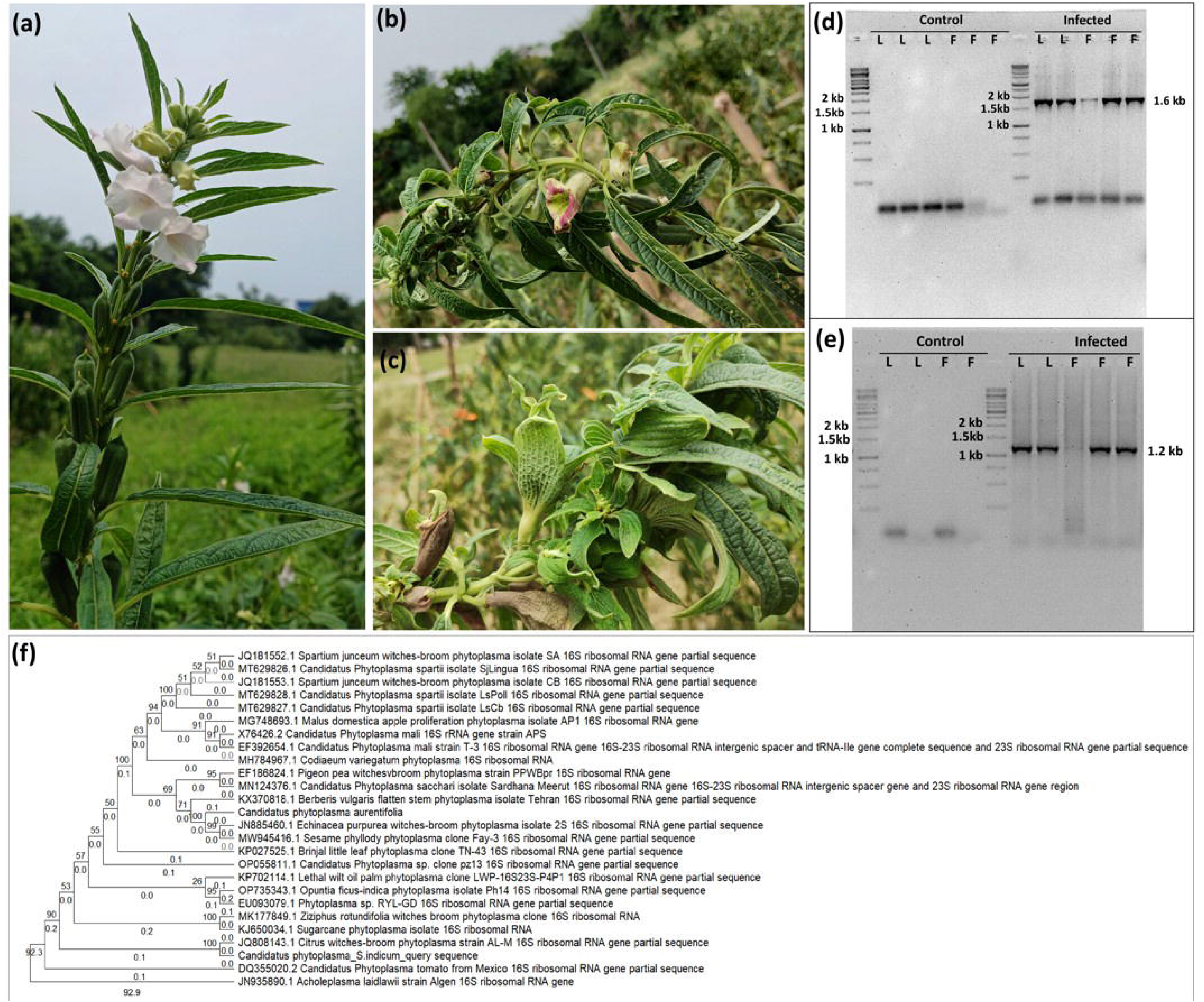
(a) Control and (b-c) symptomatic Sesame plants; (c) PCR amplification using the primer P1/P7 generated a 1.6kB band (d) Nested-PCR using R162Fn/R162R primers generated a 1.2kB band (e) Phylogenetic tree showing the detected query strain in relation to previously reported Phytoplasma sequences.

### Identification of the Phytoplasma strain and Phylogenetic analysis

Sanger sequencing of the ∼1.2 kb nested PCR amplicon, followed by phylogenetic analysis, was performed to identify the Phytoplasma strain. As shown in the bootstrapped Maximum Likelihood tree (Fig. 1f), the sequence clustered with Citrus witches’ broom Phytoplasma strain AL-M (16S rRNA gene; GenBank Accession No. JQ808143.1), exhibiting near 100% sequence identity. This AL-M strain was first reported by <u>Ghosh *et al.*, (2013)</u> as the causal agent of witches’ broom disease of acid lime (WBDL) in *Citrus aurantifolia*, and was later recognized as *Ca.* Phytoplasma *aurantifolia* by multiple authors [54].

### Comparative analyses of morphological parameters between control and infected stem

A comparative analysis between histological sections of the Phytoplasma non-infected (Fig. 2a-b) and infected sesame (Fig. c-d) stems revealed that most of the vascular tissue-associated parameters remained unaffected. For instance, no significant differences were observed in the average length of xylem vessels (control: 32.8 ± 6.8 µM; infected: 36.8 ± 9.2 µM; Mann Whitney *p* = 0.344; n=10) or the average length of vascular bundles (control: 411.3 ± 126.2 µM; infected: 460.6 ± 65.8 µM; Mann Whitney *p* = 0.851; n=10). In contrast, the cortical parenchyma cells in infected stems were marginally smaller (control: 161.9 ± 31.8 µM; infected: 135.4 ± 11.8 µM; Mann Whitney *p* = 0.052; n=10). Notably, the total cortical region length was significantly reduced in infected stems compared to controls (control: 273.6 ± 44.2 µM; infected: 114.0 ± 36.2 µM; Mann Whitney *p* < 0.0001; n=10). potentially contributing to stem flattening observed upon Phytoplasma infection [11] (Fig. 2e-j).

**Fig. 2:**
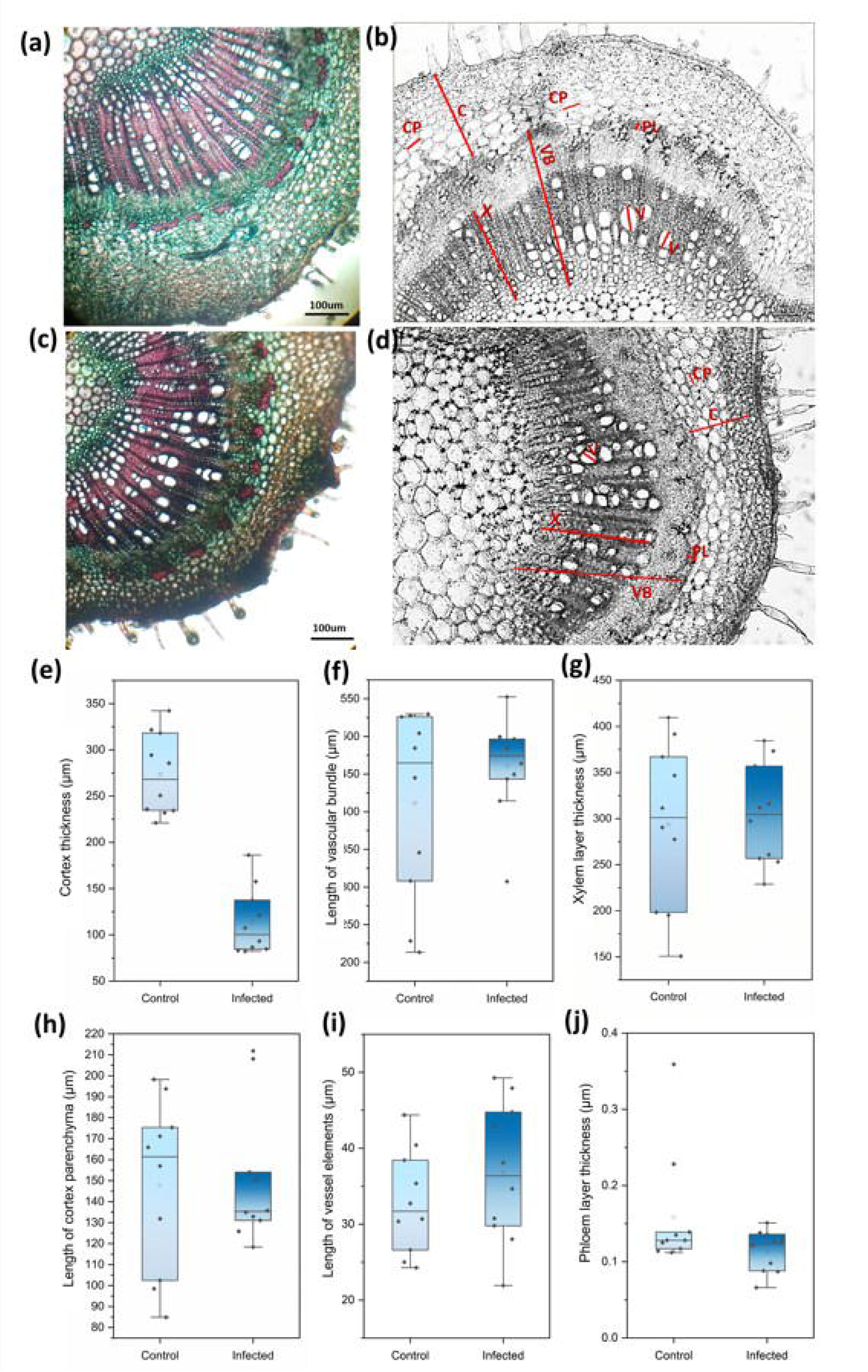
T.S of (a-b) non-infected and (c-d) infected plant stem: Study of morphological parameters, (e) Cortex thickness (C, µm), (f) Length of vascular bundle (VB, µm), (g) Xylem layer thickness (X, µm), (h) Length of cortex parenchyma (CP, µm), (i) Length of vessel elements (V, µm), (j) Phloem layer thickness(PL, µm).

## Metabolomics data Analysis

Untargeted metabolomic profiling of healthy and Ca. Phytoplasma–infected tissues (leaf and flower) revealed significant alterations in the metabolic landscape of Sesamum indicum. A total of 413 metabolites were identified, among which the most abundant classes included phenylpropanoids (31), porphyrin metabolites (31), fatty acids and derivatives (31), amino acids (31), phytohormones and their derivatives (31), among others. Many of these metabolites were differentially accumulated (DAMs) in response to infection. In leaf tissues, 227 metabolites were differentially accumulated, of which 181 were significantly over-accumulated (log2FC > 1) and 46 were significantly depleted (log2FC <-1) (Table S3). Similarly, in floral tissues, 163 metabolites were over-accumulated and 90 were depleted upon infection (Fig. 3a-c).

**Fig. 3:**
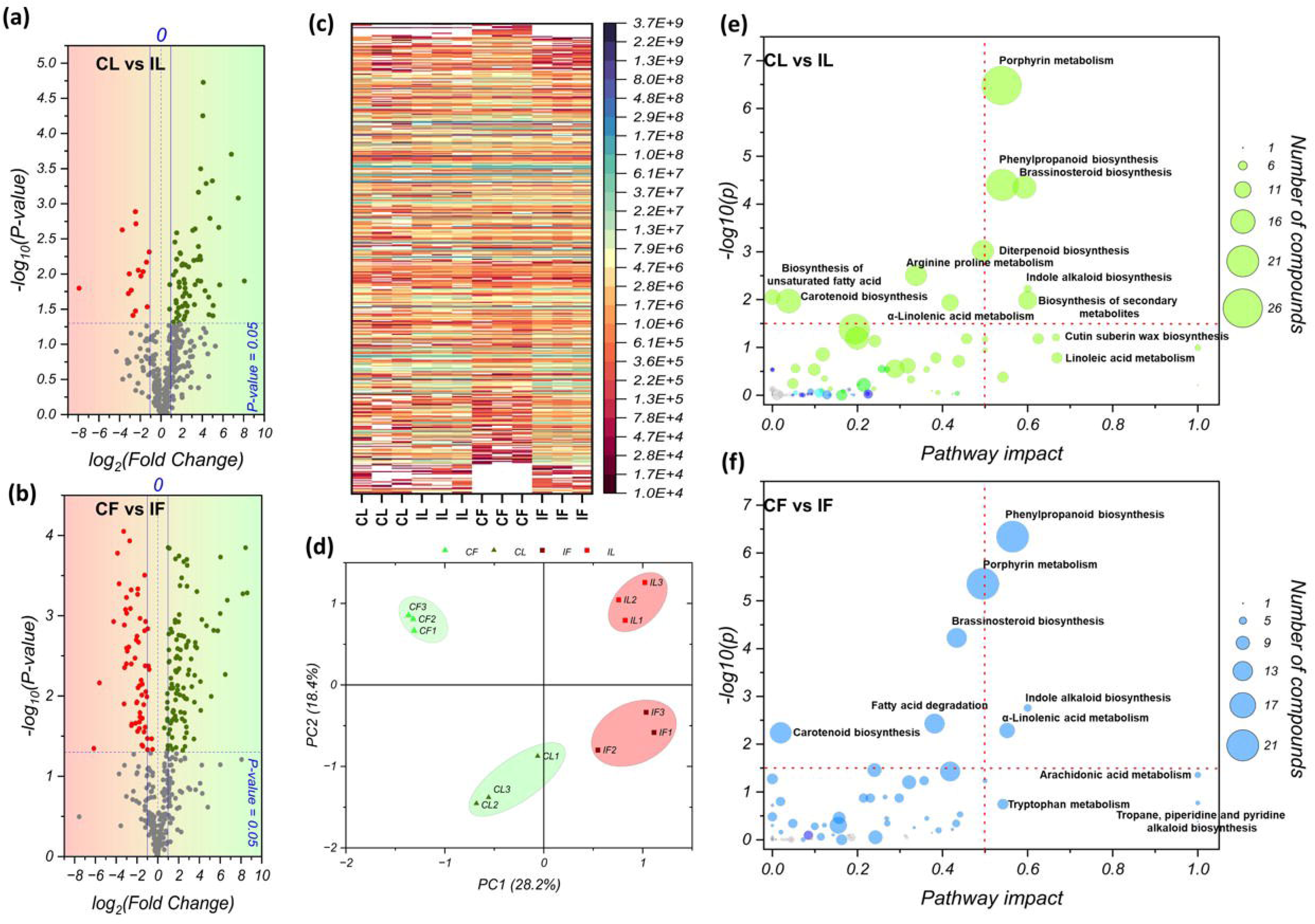
Volcano plot showing the differential accumulation of metabolites in (a) leaf and (b) flower tissues; (c) Heatmap showing the raw intensity values of all detected metabolites (y-axis) across the investigated samples (x-axis); (d) Principal component analysis (PCA) score plot illustrating the clustering of samples according to their metabolomic profiles; Pathway impact analysis plots based on differentially accumulated metabolites (DAMs) in (e) Leaf and (f) Flower, showing the key pathways that get affected upon infection. [CL – Control leaf, IL – Infected leaf, CF – Control flower, IF – Infected flower]

To examine the separation of samples based on their metabolomic profiles, principal component analysis (PCA) was performed. The PCA score plot demonstrated a clear segregation between healthy (CL and CF) and infected tissues (IL and IF), indicating distinct metabolic profiles. PC1 and PC2 together explained 46.6% of the total variance, with PC1 accounting for 28.2% (eigenvalue: 116.86) and PC2 accounting for 18.4% (eigenvalue: 76.33). The segregation of healthy and infected samples was primarily observed along PC1 (Fig. 3d).

Pathway impact analysis was further conducted using KEGG orthology-based annotations against the Arabidopsis thaliana reference metabolome dataset, which provided functional insights into the biological pathways that get affected during infection. The analysis revealed that several critical pathways, including porphyrin metabolism, phenylpropanoid biosynthesis, brassinosteroid biosynthesis, diterpenoid biosynthesis, cutin suberin wax biosynthesis, and α-linolenic acid metabolism, were significantly impacted in infected tissues. Notably, some of which were tissue-specific, indicating that Phytoplasma infection triggers a complex and compartmentalized reprogramming of host metabolism (Fig. 3e-f).

### Illumina-Seq data analysis

*De novo* transcriptome assembly of the four sample groups, i.e., control flower (CF), control leaf (CL), infected flower (IF), and infected leaf (IL), yielded a total of 101,472 unigenes. Subsequent analysis identified 76,081 unique sequences with potential protein-coding capacity. Of these, 73,498 unigenes were successfully annotated against the NCBI non-redundant (NR) protein database using BLASTX. The majority of BLAST hits (54,412 CDS) showed highest similarity to the concerned organism, i.e., *Sesamum indicum*. A smaller proportion matched other species, including *Thrips palmi* (2,835 CDS), *Paulownia fortunei* (1,639 CDS), and *Apolygus lucorum* (1,183 CDS), representing a negligible fraction of the total CDS (Figure S1). The number of CDS detected across samples, i.e., CF (28,061 – 33,397 CDS), CL (26,544 – 30,834 CDS), IF (29,912 – 31,429 CDS), and IL (27,036 – 31,206 CDS), remained within comparable ranges, indicating consistency in sequencing, assembly, and downstream analytical workflows (Table S4).

Differential expression analysis revealed distinct transcriptional responses between infected and healthy tissues (Fig. 4a). In the CF vs IF comparison, 5,086 genes were significantly upregulated (log_2_FC > 1) and 5,233 genes were significantly downregulated (log_2_FC < −1) in infected flowers relative to controls. In the CL vs IL comparison, 5,086 genes were significantly upregulated (log_2_FC > 1), while 8,236 genes were significantly downregulated (log_2_FC < −1) in infected leaves relative to controls. Tissue-specific expression analysis identified 8,207 CDS exclusively expressed in floral tissues and 4,143 CDS exclusively in foliar tissues. Additionally, 7,201 CDS were uniquely expressed in healthy tissues (leaf and/or flower), while 5,088 CDS were exclusive to infected tissues (Fig. 4b-c).

**Fig. 4:**
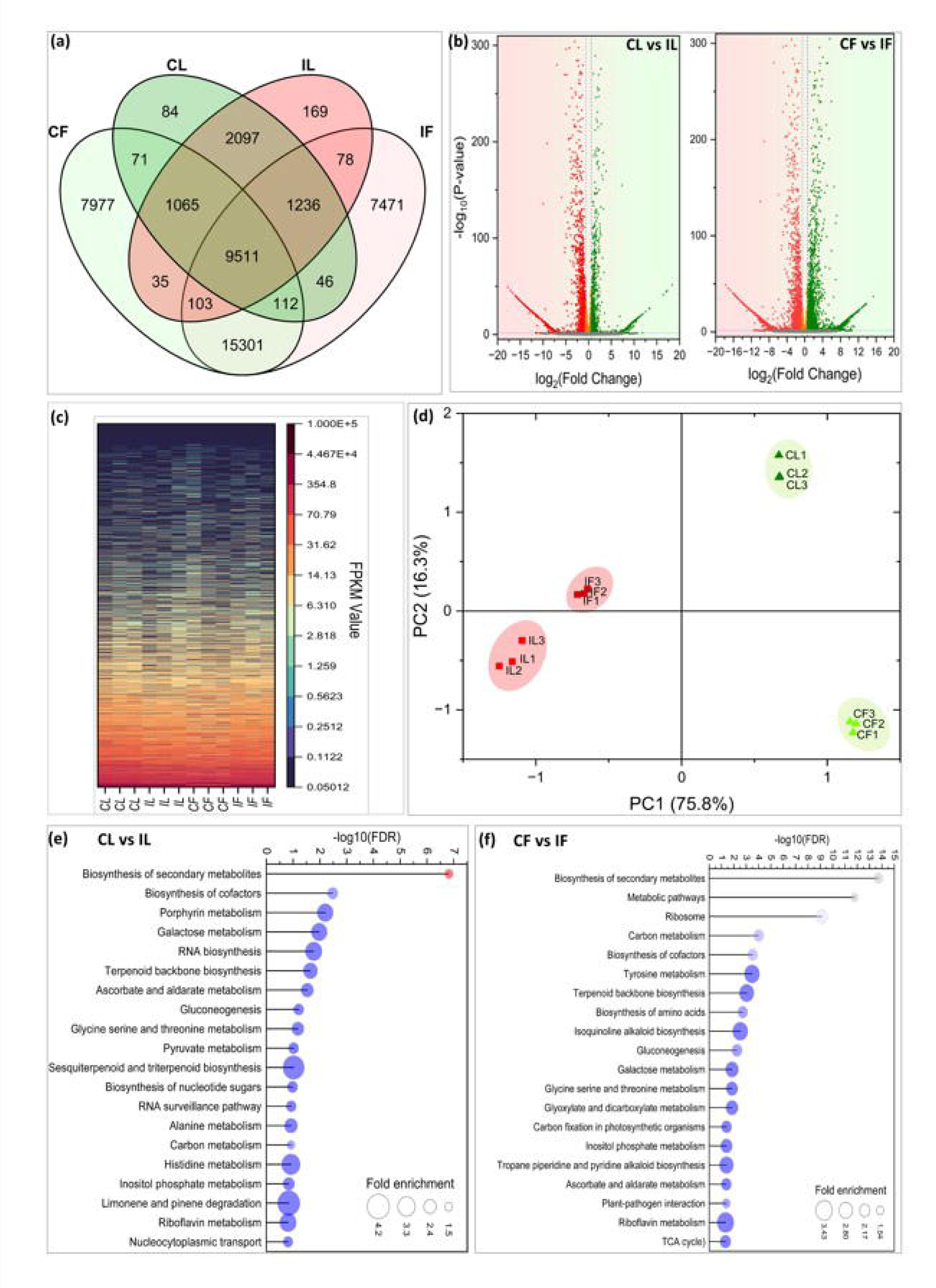
(a) Venn diagram showing the number of unique and shared CDS among the four samples. (b) Volcano plots displaying the distribution of differentially expressed genes (DEGs) in (i) leaf and (ii) flower tissues. (c) Heatmap of FPKM values for all annotated coding sequences (CDS) across the investigated samples (y-axis: genes; x-axis: samples). (d) Principal component analysis (PCA) score plot illustrating the clustering of samples based on transcriptomic profiles. (e–f) Enrichment analysis plots based DEGs in (e) leaf and (f) flower tissues, representing key pathways that get affected upon infection. [CL – Control leaf, IL – Infected leaf, CF – Control flower, IF – Infected flower]

Principal component analysis (PCA) demonstrated clear separation between infected and healthy tissues. PC1 accounted for 75.8% of the total variance, while PC2 explained 16.3%. Infected samples (IL and IF) clustered distinctly on the negative axis of PC1, whereas healthy samples (CF and CL) clustered on the positive axis. The tight clustering of biological replicates within each group further confirmed the reproducibility and reliability of the transcriptomic data. Notably, based on the overall gene expression profiles, infected tissues (both floral and foliar) exhibited greater similarity to each other than to their respective healthy counterparts, as evidenced by their close clustering on the negative axis of PC1, while control samples clustered on the positive axis (Fig. 4d).

KEGG pathway enrichment analysis of DEGs revealed that, in CF vs IF, significantly enriched pathways included biosynthesis of secondary metabolites, tyrosine metabolism, terpenoid backbone biosynthesis, alkaloid biosynthesis (isoquinoline, tropane, piperidine, and pyridine), ascorbate metabolism, and plant - pathogen interactions. In CL vs IL, enriched pathways predominantly included biosynthesis of secondary metabolites, porphyrin metabolism, terpenoid backbone biosynthesis, ascorbate metabolism, and sesquiterpenoid–triterpenoid biosynthesis (Table S5, Fig. 4e-f).

### Disruption of Flowering Gene Networks and Floral pigment Biosynthesis

As Phytoplasma infection resulted in the development of floral virescence and phyllody, we decided to explore the expression patterns of genes that regulate flowering. Transcriptomic analyses of control and infected flowers revealed several differentially expressed genes (DEGs) associated with flowering mechanisms in angiosperms, exhibiting dynamic expression patterns. All the identified major genes involved in floral organ identity development, such as *apetala* (AP), *sepallata* (SEP) and *agamous* (AG) exhibited down regulating trends, especially *apetala*-2 (log_2_FC:-1.68) and *sepallata*-X2 (log_2_FC:-1.50) exhibited significant downregulation upon infection (Fig. 5a). Other genes involved in Petal and stamen identity (*deficiens* and *globosa*) organ identity, such as *deficiens-like* (log_2_FC:-3.54), *globosa* (log_2_FC:-1.88) and *globosa-like* (log_2_FC:-1.68) also exhibited strong downregulation. On the other hand, genes involved in tissue meristem identity such as *squamosa promoter binding protein*, *clavata* and *wuschel* mostly exhibited upregulating trend. For example, SPL6 (log_2_FC: 1.44), *SPL9* (log_2_FC: 1.35), *SPL16* (log_2_FC: 1.19) *clavata* 3 (log_2_FC: 1.79) exhibited strong upregulation pattern (log_2_FC: 1.79). A few genes belonging to these groups such as *clavata*1, *wuschel* related homeobox 8, *wuschel* related homeobox 4 and its two other isoforms X1 and X3 were exclusive detected in significantly high FPKM in Phytoplasma-infected sesame flower. Other genes associated with flowering, that governs flowering time or flowering pattern, e.g*., FLD, FLK* and *constans*, did not exhibit specific uniform upregulating or downregulating trend. For example, some transcripts of *constans* exhibit significant upregulation while some other exhibit strong down regulation.

**Fig. 5:**
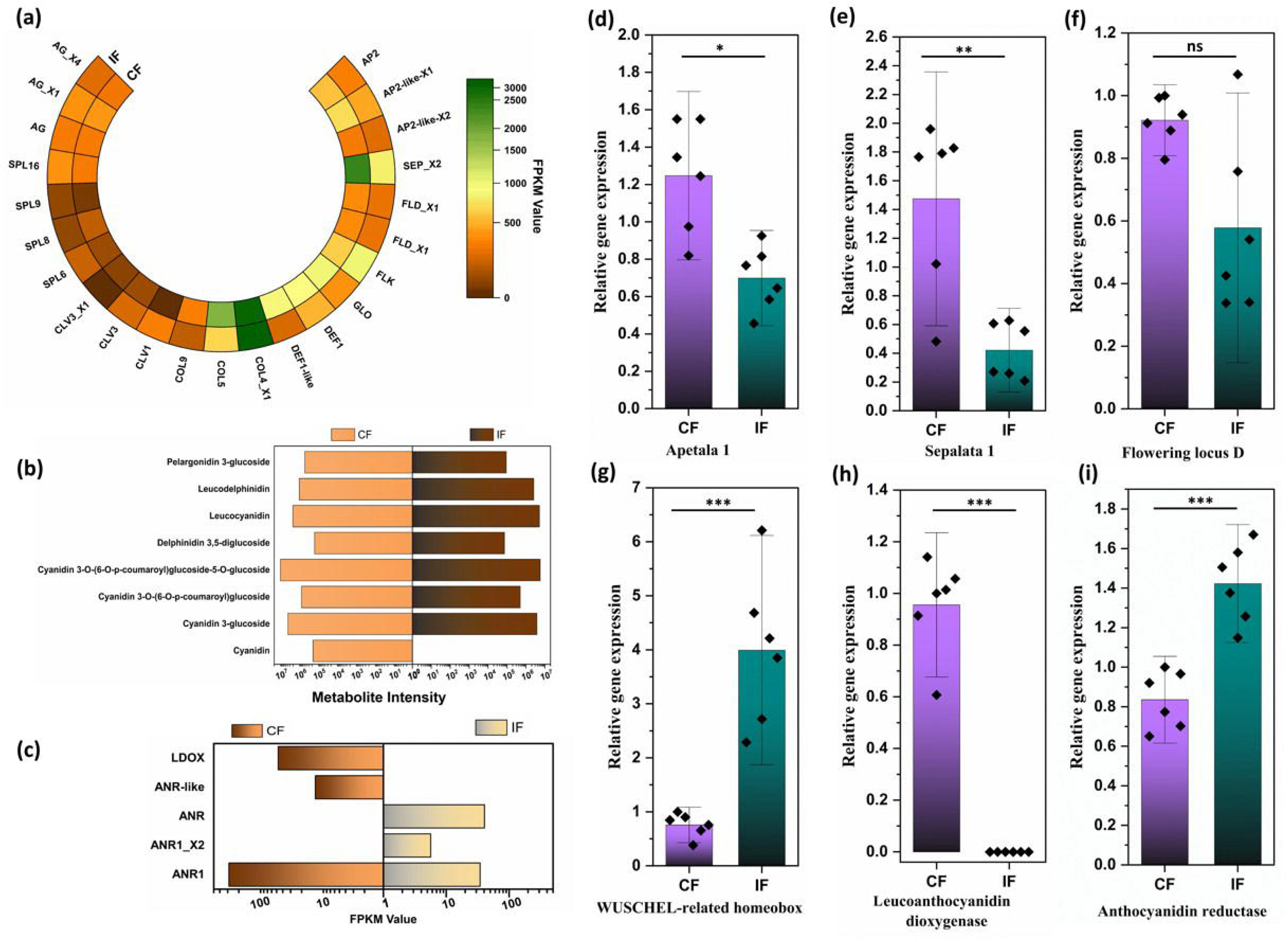
(a) Heatmap showing FPKM values of genes associated with flower development. Comparative bar diagrams depicting (b) metabolite abundance and (c) FPKM values of genes involved in the anthocyanin biosynthetic pathway. Gene expression validation by qRT-PCR of (d) APETALA1 (AP), (e) SEPALLATA1 (SEP), (f) FLOWERING LOCUS D (FLD), (g) WUSCHEL-related homeobox, (h) LEUCOANTHOCYANIDIN DIOXYGENASE (LDOX), and (i) ANTHOCYANIDIN REDUCTASE (ANR).

Another floral-specific gene alteration was observed in the pigment biosynthetic pathway. The coloration of sesame flowers is primarily determined by anthocyanins, and Phytoplasma infection markedly suppressed the expression of *anthocyanidin 3-O-glucosyltransferase*, *anthocyanidin 3-O-glucoside 6’’-O-acyltransferase-like*, and *leucoanthocyanidin dioxygenase* (log2FC <-1) (Fig. 5b). At the metabolite level, cyanidin, the pigment responsible for the characteristic slight purple colour of sesame flowers, was detected exclusively in healthy floral tissues and was completely absent in infected samples. Several cyanidin glucosides, including cyanidin 3-O-β-D-sambubioside and cyanidin 3-O-(6-O-p-coumaroyl)glucoside, also showed a reduction in abundance, though the changes were not statistically significant. In contrast, the precursor compounds, leucoanthocyanidins, accumulated significantly (e.g., leucocyanidin, log2FC = 1.33), suggesting inefficient or impaired conversion of leucoanthocyanidins into anthocyanins in infected flowers (Fig. 5c). The expression of few genes, namely, *AP1*, *SEP1, FLD*, *WUSHCHEL*-related homeobox, *Leucocyanidin dioxygenase (LDOX*) and *Anthocyanidin reductase (ANR*), were also validated using qRT-PCR (Fig. 5d-i).

### Phytoplasma causes tissue specific alteration of chlorophyll pathway

Among the phenotypic alterations observed upon Phytoplasma infection in sesame plants, tissue-specific changes in green pigmentation, or virescence, is one of the most prominent characters (Fig. 6a). Quantification of photosynthetic pigments using UV-Vis spectrophotometry revealed a substantial increase (*t=*14.49; *p=*0.011) in total chlorophyll content in floral tissues of infected plants (49.15 ± 4.67 mg/g), compared to the control plants (5.38 ± 1.42 mg/g). Similarly, foliar tissues of infected plants also showed an elevated chlorophyll content (56.75 ± 3.82 mg/g) relative to control leaves (50.81 ± 6.86 mg/g), although the increase was not statistically significant (Fig. 6 b-c; Table S6).

**Fig. 6:**
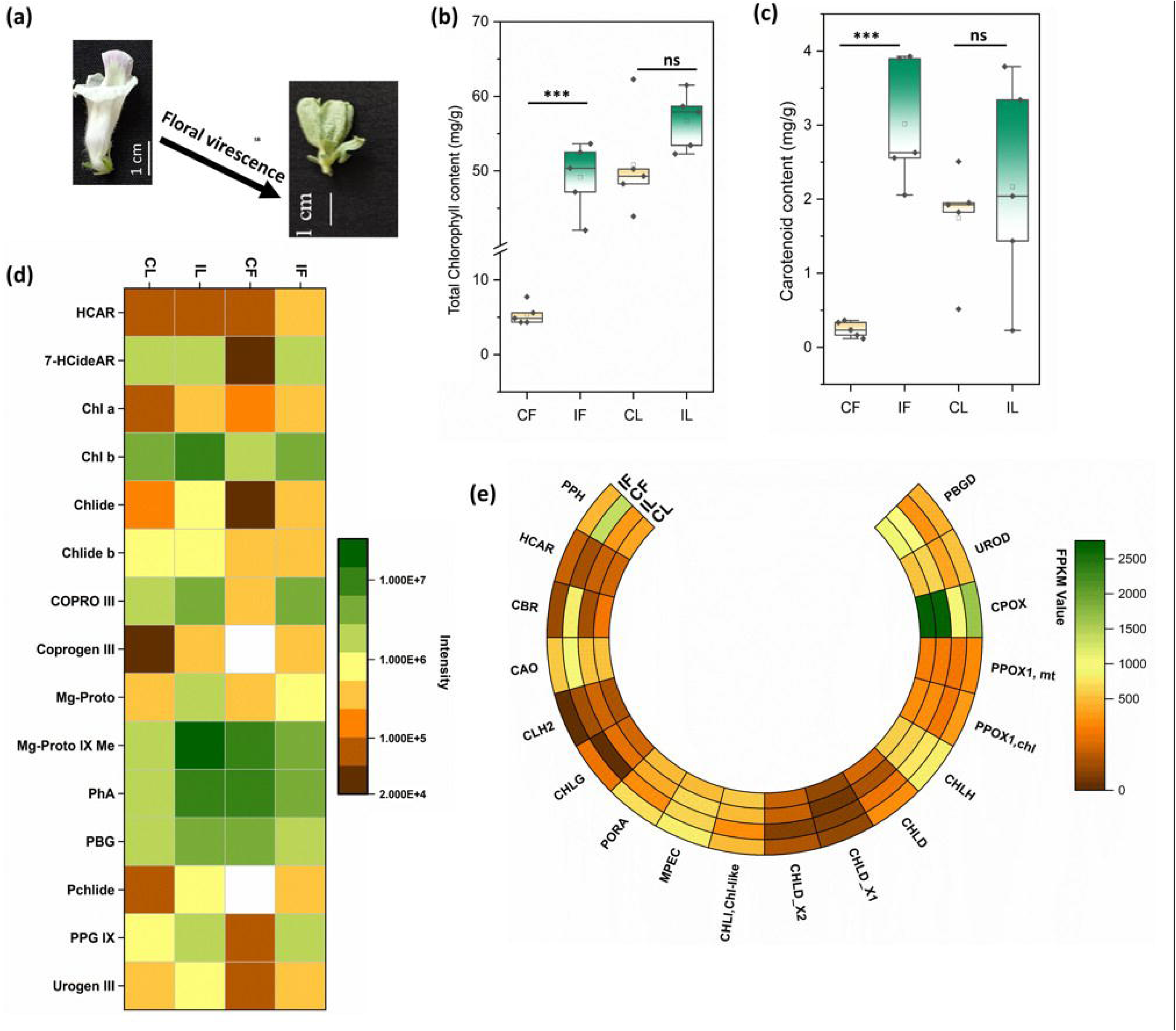
(a) Phytoplasma-induced development of green pigmentation in floral tissues (floral virescence). Spectrophotometric quantification of (b) total chlorophyll content (mg/g) and (c) carotenoid content (mg/g). (d) Heatmap showing the abundance of metabolites and (e) FPKM values of genes associated with chlorophyll (porphyrin) biosynthesis and degradation.

Metabolomic profiling using LC-MS/MS further elucidated the differential abundance of key metabolites involved in the porphyrin and chlorophyll biosynthesis pathway across infected and non-infected tissues. In foliar tissues, Phytoplasma infection led to significant upregulation of several intermediates, including uroporphyrinogen III (log_FC: 1.73), coproporphyrinogen III (log_FC: 3.62), coproporphyrin III (log_FC: 1.74), magnesium protoporphyrin (log_FC: 1.74), magnesium protoporphyrin monomethyl ester (log_FC: 4.26), protochlorophyllide (log_FC: 3.17), chlorophyllide (log_FC: 2.71), and chlorophyll a (log_FC: 2.60). Other metabolites such as Porphobilinogen (log_FC: 0.87) and chlorophyll b (log_FC: 0.95) also exhibited enhanced accumulation.

The infected floral samples showed a marked increase in chlorophyll a (log_FC: 1.15) concerning the control flowers, while chlorophyll b also exhibited a non-significant, increase (log_FC: 0.95). The key intermediates such as protoporphyrinogen IX (log_FC: 4.80), Coproporphyrin III (log_FC: 4.09), uroporphyrinogen III (log_FC: 2.06), 7(1)-hydroxychlorophyll a (log_FC: 1.90), chlorophyllide (log_FC: 2.56), and 7(1)-hydroxychlorophyllide (log_FC: 6.06) accumulated significantly in infected floral tissues (Fig. 6d).

Transcriptome analysis supported these metabolic findings, with genes involved in chlorophyll biosynthesis showing upregulation in infected tissues. In foliar tissues, genes such as *porphobilinogen deaminase* (log_FC: 0.135), *uroporphyrinogen decarboxylase* (log_FC: 0.203), *magnesium-protoporphyrin IX monomethyl ester oxidative cyclase* (log_FC: 0.47), and *chlorophyll synthase* (log_FC: 0.44) exhibited increasing trends, though not reaching statistical significance. In contrast, floral tissues displayed more pronounced transcriptional responses. Genes such as *magnesium-chelatase subunit II* and *protochlorophyllide reductase* were significantly upregulated (log_FC: 1.77 each), while *chlorophyll synthase* was detected in a significantly higher quantity (FPKM: 130.83) exclusively in infected floral tissues. Other biosynthetic genes, including *porphobilinogen deaminase* (log_FC: 0.84), *uroporphyrinogen decarboxylase* (log_FC: 0.59), and *coproporphyrinogen iii oxidase 1* (log_FC: 0.86), also showed an upregulating trend (Fig. 6e).

Given that chlorophyll levels are regulated by both synthesis and degradation, metabolites involved in chlorophyll catabolism were also analyzed. In infected foliar tissues, catabolites such as Pheophorbide a (log_FC: 2.22), Primary Fluorescent Chlorophyll Catabolite (log_FC: 1.55), and Red Fluorescent Catabolites (log_FC: 2.036) showed significant accumulation. Conversely, in floral tissues, Pheophorbide a exhibited a downregulation trend. While both Red Chlorophyll Catabolite (log_FC: 1.45) and Primary Fluorescent Chlorophyll Catabolite (log_FC: 4.012) were significantly downregulated in infected flowers. These changes were also reflected at the transcript level. In foliar tissues, chlorophyll degradation-related genes such as *chlorophyllase* and *pheophytinase* (log_FC: 1.3) were upregulated. In contrast, in floral tissues, both *chlorophyllase* and *pheophytinase* were significantly downregulated (log_FC:-2.031), consistent with the suppressed accumulation of degradation products.

### Phytoplasma infection changes ROS metabolism

As the generation of oxidative stress is one of the inevitable phenomena that occur as a consequence of infection, ultimately triggering various downstream pathways, we analyzed different antioxidant scavenging activities and stress enzyme activities using UV-Vis spectrophotometry. The study revealed that the total antioxidant activity (TAA) increased in the infected leaves and flowers upon Phytoplasma infection compared to the healthy ones. DPPH and ABTS radical scavenging activities were higher in both floral and foliar tissues following Phytoplasma infection. The FRAP assay, commonly used to measure antioxidant capacity, also showed increased activity in both infected flower and leaf samples compared to the healthy controls. Furthermore, superoxide (SO) radical scavenging activity was considerably higher in the infected leaf and floral samples, indicating a strong capacity of the samples to neutralize these harmful radicals (Fig. 7a-e).

**Fig. 7:**
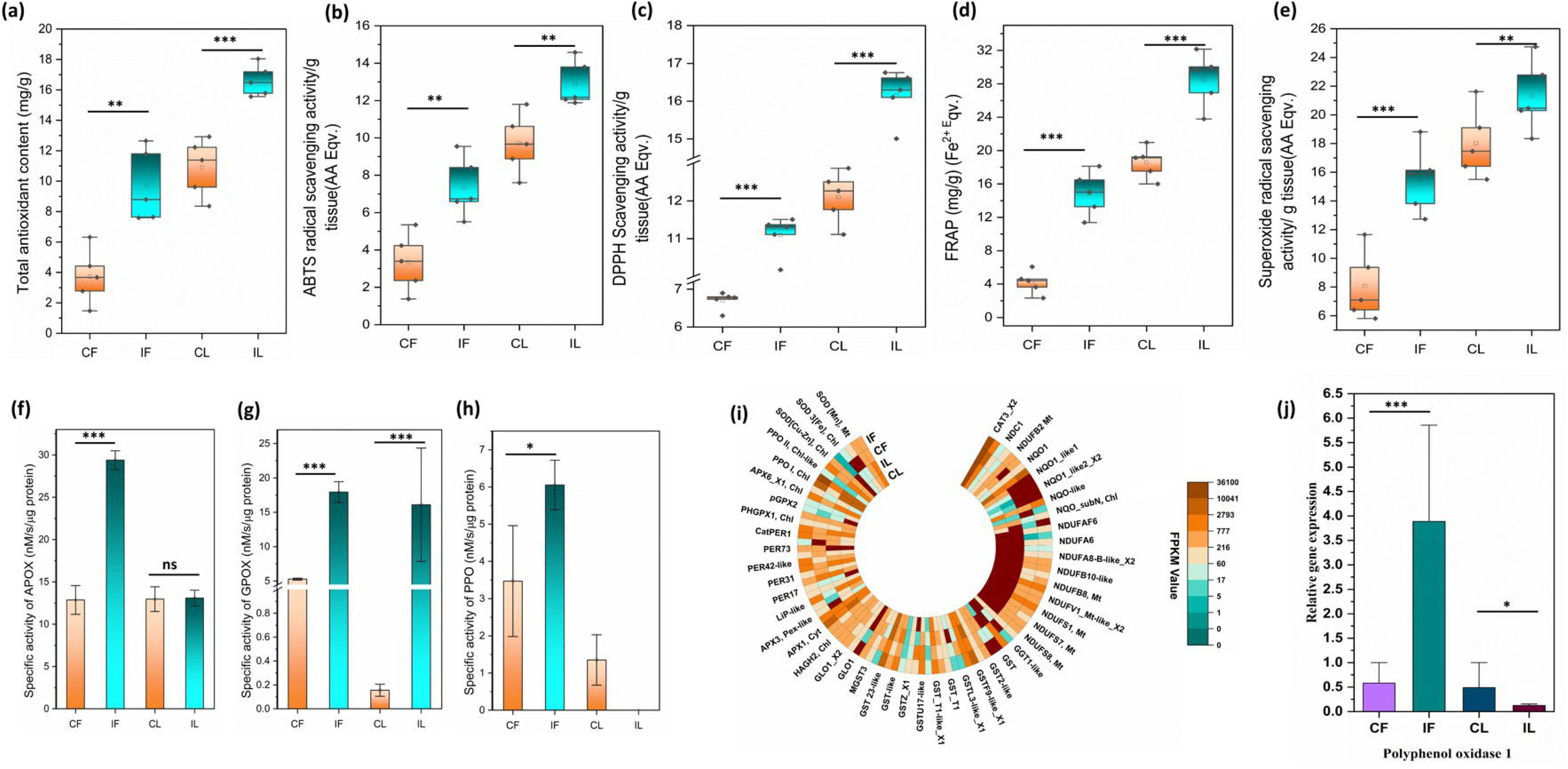
Box plots representing (a) total antioxidant capacity, (b) ABTS, (c) DPPH, (d) FRAP, and (e) superoxide (SO) scavenging activity in the investigated tissues before and after infection. Bar diagram showing the specific activity of (f) APOX, (g) GPOX, and (h) PPO in the tested samples. (i) Heatmap showing FPKM values of CDS associated with oxidative stress across all samples. (j) Relative gene expression of Polyphenol oxidase (PPO1) in the tested samples, validated by qRT-PCR.

Next, we decided to look over the stress or antioxidant enzymes that help in neutralizing the oxidative damage caused by ROS. Among the scavenging enzymes, the specific activity of Guaiacol peroxidase (GPOX) increased upon infection in both the floral and foliar tissues when compared to their control samples. However, considerable tissue specific variation in GPOX activity was noted in the studied tissues (Fig. 7f). Another stress enzyme like Ascorbate peroxidase (APOX) activity also increased in the infected floral samples than the healthy ones, whereas APOX activity remains non-significantly altered in the infected leaves (Fig. 7g). The activity of another enzyme Polyphenol Oxidase I (PPO I), significantly increased in floral tissues upon infection, although, its activity was almost negligible in the infected leaves when compared to the control leaves (Fig. 7h).

These physiological findings were corroborated by transcriptomic data, which demonstrated variations in the expression levels of linked genes, upon Phytoplasma infection. The genes involved in biosynthesis of stress enzymes or mitigating oxidative stress exhibited differential expression pattern upon infection in the transcriptomics analysis. Similar to antioxidant data, obtained from UV-Vis spectrophotometry analysis, the data also show that there are considerable tissue specific variation between responsiveness towards infection (in compared to control). The early stress-responsive gene class, *SOD*, displayed an overall upregulation trend in floral tissues. In floral tissues, infection induced significant upregulation of *superoxide dismutase* [Mn] (log_FC = 1.18) and *superoxide dismutase* [Cu-Zn]-like (log_FC = 1.18). Additionally, *superoxide dismutase* [Fe], chloroplastic, and *superoxide dismutase* [Fe] 3, chloroplastic, exhibited an exclusive expression pattern in response to infection in floral tissues. In leaf tissues, members of the *SOD* gene family displayed variable expression patterns, with some showing upregulation and others downregulation; however, none of the changes were statistically significant. Another early stress-responsive gene*, catalase* (CAT), did not show notable differential expression in either tissue type. The *peroxidase* gene family represented one of the most abundant classes of stress-responsive genes, exhibiting significant differential expression across tissue types upon infection. In leaf tissues, a total of 23 *peroxidase* genes were detected, of which eight members were significantly upregulated (log_FC > 1). Notable examples include *cationic peroxidase 1* (log_FC = 2.15), *probable L-ascorbate peroxidase 6*, chloroplastic isoform X2 (log_FC = 1.04), and *L-ascorbate peroxidase 3*, peroxisomal-like (log_FC = 1.02). While several members displayed downregulation, none reached the significance threshold (log_FC < –1). In floral tissues, the expression profile of *peroxidase* genes was more dynamic. A total of 33 *peroxidase* genes were detected, among which nine were significantly upregulated and three significantly downregulated upon infection. Noteworthy members included *lignin-forming anionic peroxidase-like* (log_FC = 4.26), *lignin-forming anionic peroxidase* (log_FC = 4.61), *probable L-ascorbate peroxidase 6*, chloroplastic isoform X1 (log_FC = 1.62), and *cationic peroxidase 2-like* isoform X2 (log_FC = 1.92). In contrast, *peroxidase 31* (log_FC = –1.55), *peroxidase 47* (log_FC = –1.29), and *peroxidase 6* (log_FC = –7.06) were markedly downregulated. Additionally, five *peroxidase* genes were exclusively upregulated and six exclusively downregulated in floral tissues upon infection (Fig. 7 i).

Glutathione-associated genes, which play a pivotal role in ROS detoxification, also showed tissue-specific regulation. In leaf tissues, 38 genes associated with glutathione-mediated ROS quenching were detected. Within this group, *glutathione S-transferase* (GST) genes exhibited a mixed expression pattern, with seven members being upregulated or uniquely expressed upon infection, while five were downregulated. In floral tissues, 45 glutathione-related genes were identified, including significant upregulation of *glutathione synthases* (2 genes) and *glutathione reductase* (1 gene). GST genes again showed variable expression, with eight members upregulated and seven downregulated following infection.

*Polyphenol oxidase* (PPO) genes exhibited a contrasting expression pattern between tissues. In infected leaves, *PPO I* (log_FC = –0.073) and *PPO II* (log_FC = –2.489) showed reduced expression, whereas in infected floral tissues, *PPO I* (log_FC = 5.42) and *PPO II* (log_FC = 2.32) were strongly induced, corroborating the data of spectrophotometric enzyme activity and qRT-PCR (Fig. 7j).

### Integrated Transcriptomic and Metabolomic parameters of Octadecanoid Pathway

The octadecanoid (lipoxygenase) pathway is a key defense signalling cascade that utilizes α-linolenic acid as its primary substrate. Through a series of intermediate reactions, this pathway ultimately produces jasmonic acid, a major phytohormone central to plant defense regulation (Fig. 8a). Similar to oxidative stress responses, the octadecanoid pathway exhibited distinct tissue-specific metabolic profiles In leaf tissues, early pathway metabolites such as α-linolenic acid, 13(S)-HPOT, OPC4-CoA, jasmonic acid (Log_FC: 0.28), and its derivative methyl jasmonate (Log_FC: 0.51) displayed a trend of increased accumulation upon infection, although in most cases the changes did not surpass the threshold for significant differential expression (Log_FC > 1) (Fig. 8b). In contrast, floral tissues demonstrated a markedly different pattern, with α-linolenic acid, OPC4-CoA, OPC6-CoA, and trans-2-enoyl-OPC8-CoA being either significantly upregulated (Log_FC > 1) or uniquely detected following infection (Fig. 8c). Jasmonic acid also accumulated to a greater extent in infected floral tissues (Log_FC: 0.51) compared to infected leaves. Notably, methyl jasmonate, which exhibited enhanced accumulation in infected leaves (Log_FC: 0.51), was significantly downregulated in infected floral tissues (Log_FC:-1.17), highlighting a key divergence in defense signalling between tissue types.

**Fig. 8:**
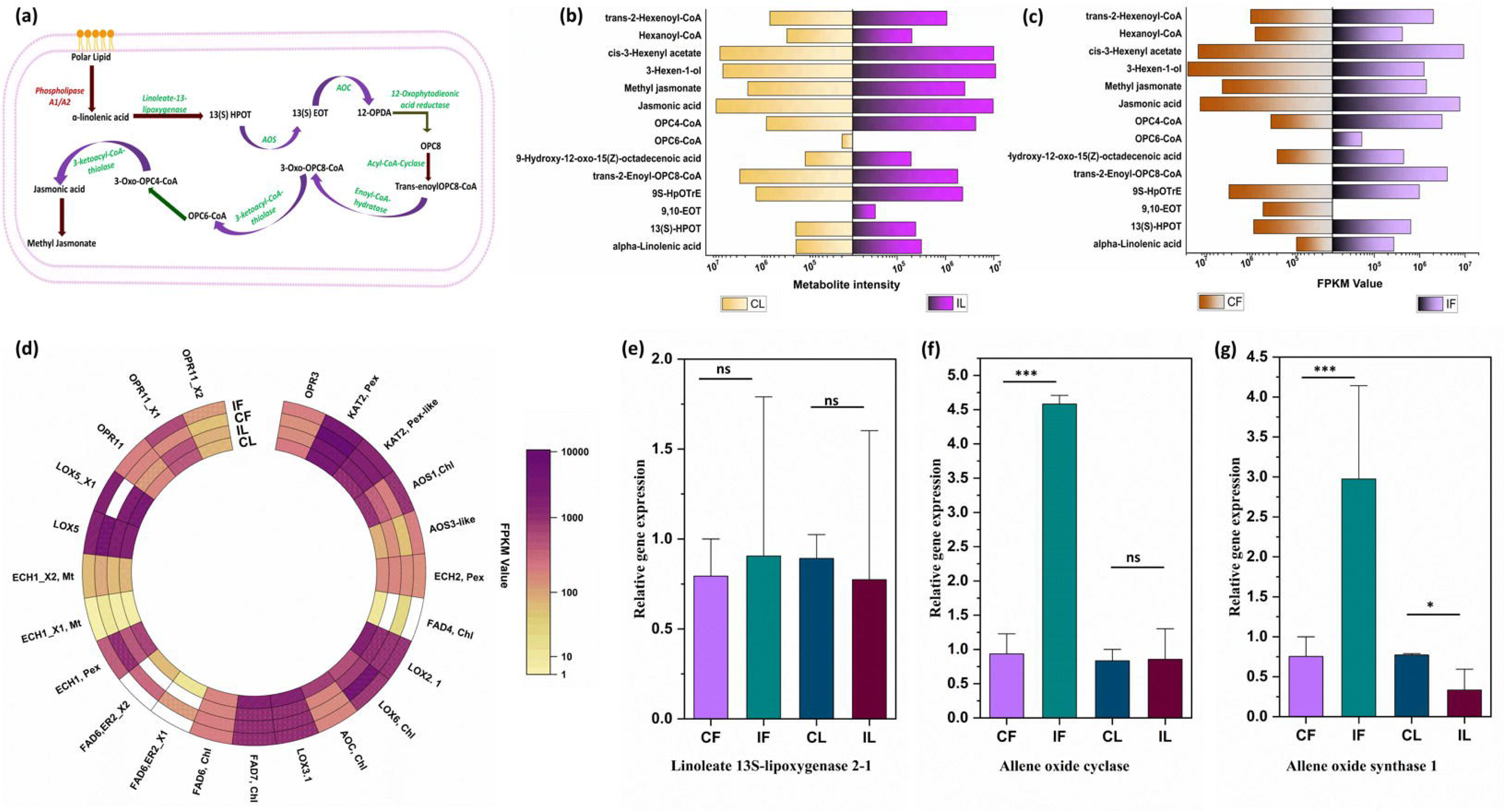
(a) (a) Diagrammatic representation of the octadecanoid pathway, with genes identified in our dataset shown in green and those not detected shown in red. (b) Metabolite accumulation in leaf and floral tissues. (c) Heatmap showing the expression of each CDS (FPKM values) associated with the octadecanoid pathway. qRT-PCR validation of (d) Linoleate 13S-lipoxygenase 2-1, (e) Allene oxide cyclase (AOC), and (f) Allene oxide synthase 1 (AOS1). [CL – Control leaf, IL – Infected leaf, CF – Control flower, IF – Infected flower].

The transcriptome data aligned with these metabolomic observations to certain extent. In foliar tissues, most genes encoding key enzymes in the pathway, such as linoleate 13S-lipoxygenase and omega-3 fatty acid desaturases, exhibited relatively stable expression patterns upon infection. Two allene oxide synthase (AOS) genes were detected, one showing an upregulation trend and the other a downregulation trend. Importantly, allene oxide cyclase (AOC), a pivotal enzyme in jasmonic acid biosynthesis, was significantly downregulated in infected foliar tissues (Log_FC:-1.77), which could result in the lack of significant changes in jasmonic acid accumulation in leaves upon infection.

In floral tissues, infection had a more pronounced impact on transcriptomic regulation of the octadecanoid pathway. Genes involved in the early chloroplast-localized steps, such as probable allene oxide synthase 3-like, allene oxide synthase 1 (chloroplastic), and two isoforms of putative 12-oxophytodienoate reductase 11 (X1 and X2), were significantly upregulated (Log_FC > 1). In contrast, several peroxisomal genes associated with the later stages of the pathway, including probable enoyl-CoA hydratase 1 (peroxisomal) and probable enoyl-CoA hydratase 2 (mitochondrial isoform X1), were significantly downregulated (Log_FC <-1), suggesting compartment-specific modulation of pathway activity (Fig. 8d).

Furthermore, another group of genes encoding linoleate 9S-lipoxygenases, enzymes responsible for producing “death acids” from the same α-linolenic acid precursor, exhibited heterogeneous responses to infection, with two genes significantly upregulated and two significantly downregulated. This complex transcriptional regulation highlights the spatial and functional partitioning of lipid-derived defense signalling in response to infection. Few genes of this pathway, namely, AOC, AOS and 13 LOX2-1, were also validated using qRT-PCR (Fig. 8 e-g).

### Impact of Phytoplasma infection on phenylpropanoid pathway

The present study reveals that *Ca.* Phytoplasma infection induces a significant accumulation of phenylpropanoids and their derivatives in both the examined tissues, with a more profound effect in flowers. The spectrophotometric analyses indicated considerable increases in total phenolic content, total flavonoids, and tannins in both tissues, particularly in floral samples (Fig. 9 a-c). Metabolomic profiling detected 22 metabolites associated with the phenylpropanoid pathway. In foliar tissue, five metabolites were significantly upregulated (Log_FC > 1) upon infection, among which coniferaldehyde, sinapyl alcohol, and caffeyl alcohol were notable. Two metabolites were significantly downregulated, including coniferin (Log_FC:-3.23), while 4-coumaryl alcohol was uniquely detected in infected leaves.

**Fig. 9:**
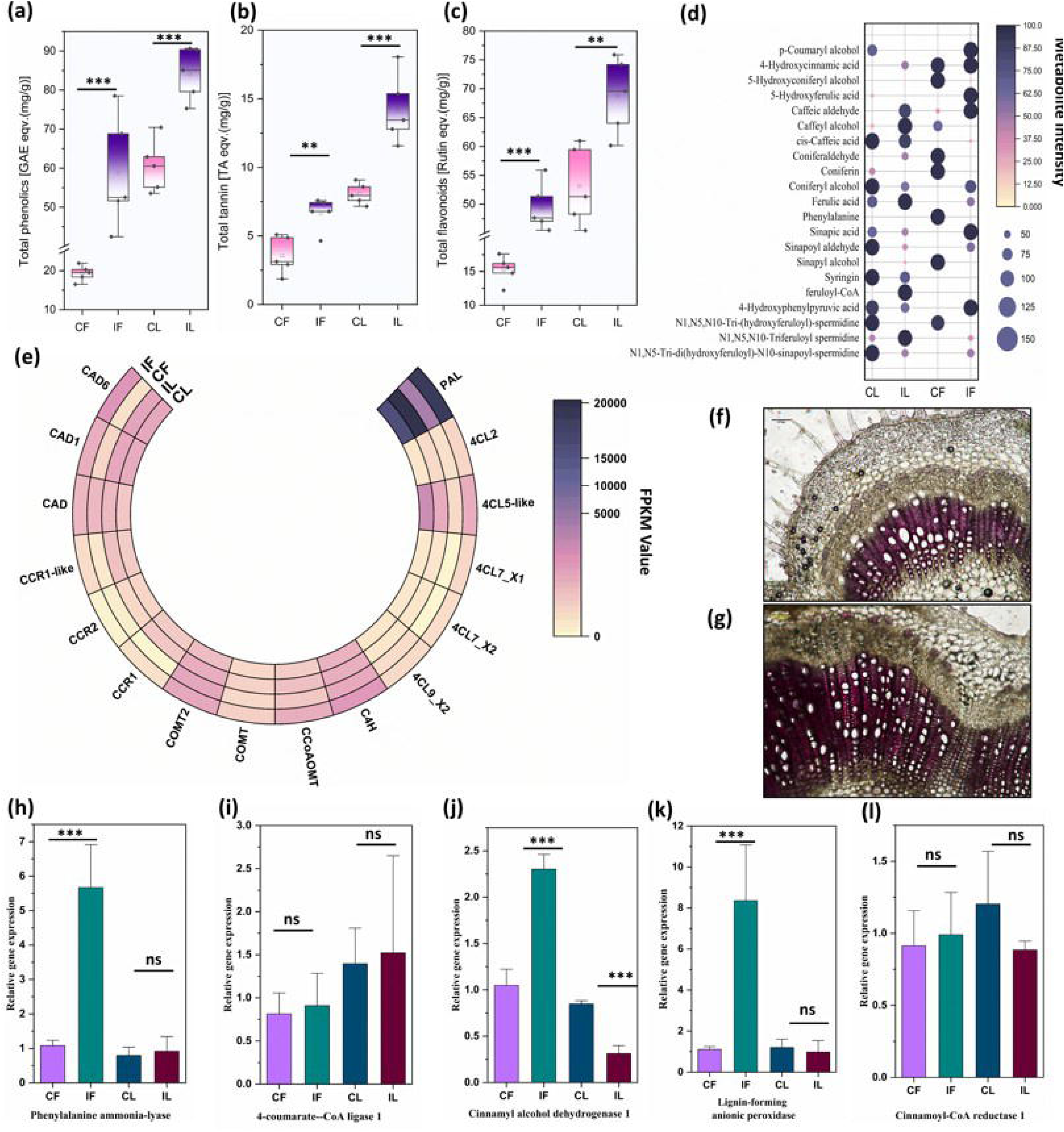
(a–c) Box-and-whisker plots showing the quantities of total phenolics, total tannins, and total flavonoids, measured using UV–Vis spectrophotometry. (d) Bubble plot illustrating the differential accumulation of metabolites in CL, IL, CF, and IF [CL – Control leaf, IL – Infected leaf, CF – Control flower, IF – Infected flower]. (e) Heatmap of the transcripts (FPKM Values) of phenylpropanoid pathway; Histochemical staining of T.S of (f) non-infected and (g) infected stem with Phloroglucinol-HCL. qRT-PCR analysis of the genes - (h) Phenylalanine-ammonia lyase (PAL), (i) 4 Coumarate-CoA ligase 1; (j) Cinnamyl alcohol dehydrogenase 1 (CAD), (k) Lignin-performing anionic peroxidase, and (l) Cinnamoyl-CoA reductase1 (CCR1).

Consistent with previous observations, the impact of Phytoplasma infection on floral tissues was more substantial in terms of modulation of phenylpropanoid pathway. Fourteen out of the 22 detected metabolites were either significantly upregulated (Log_FC > 1) or uniquely identified in infected flowers. Notable examples include 4-coumaryl alcohol, coniferyl alcohol, p-coumaric acid, caffeic acid, ferulic acid, sinapic acid, and syringin. Conversely, five metabolites, including coniferin, coniferaldehyde, and sinapyl alcohol, were significantly downregulated (Log_FC <-1) in infected floral tissues (Fig. 9d).

At the transcriptomic level, a similar trend was observed. In foliar tissues, most genes associated with the phenylpropanoid pathway showed an overall upregulation trend, including one cinnamoyl-CoA reductase (CCR) gene and one trans-cinnamate 4-monooxygenase (C4H) gene that were considerably upregulated. Only a few genes were downregulated, among which one caffeic acid 3-O-methyltransferase (COMT) gene exhibited significant downregulation. The 4-coumarate–CoA ligase (4CL) gene family displayed variable expression, with one member significantly upregulated and another significantly downregulated (Fig. 9e).

In floral tissues, consistent with the metabolomics data, most of the genes involved in the phenylpropanoid pathway, were significantly overexpressed (Log_FC > 1). Although a few genes showed downregulation, none reached statistical significance (Log_FC <-1). Genes belonging to phenylalanine ammonia lyase (PAL; 3 genes), cinnamyl alcohol dehydrogenase (CAD; 2 genes), 4CL (4 genes), CCR (2 genes), C4H (2 genes), COMT (1 gene), and caffeoyl-CoA O-methyltransferase (CCoAOMT; 1 gene) families were significantly upregulated in infected floral tissues, supporting the observed overaccumulation of phenylpropanoid metabolites in the flowers. Phloroglucinol-HCL staining of non-infected and infected stem shows higher intensity in the infected tissue, suggesting higher lignin accumulation. (Fig. 9fg) For validation using qRT-PCR, genes such as PAL, 4-coumarate-CoA ligase 1, CAD1, lignin-forming peroxidase and CCR1, were selected (Fig. 9h-l).

Lignans, a class of polyphenols, exhibited a tissue-specific differential accumulation pattern. Compounds such as pinoresinol (Log_FC=1.71) and secoisolariciresinol (Log_FC=1.47) were significantly accumulated in leaf tissues upon Phytoplasma infection, whereas they were markedly depleted in floral tissues (Log_FC=-2), strongly indicating a tissue-specific differential response in host plants.

### Phytoplasma modulates Brassinosteroid Pathway

One of the most consistent and unambiguous changes observed in response to Phytoplasma infection, in metabolomics data, regardless of whether floral or foliar tissues were examined, was the pronounced overaccumulation of both major and intermediate metabolites of the brassinosteroid biosynthesis pathway. In total, 16 brassinosteroid-related metabolites were detected in foliar tissues and 15 in floral tissues. Among these, 15 metabolites in leaves and 13 in flowers exhibited significant hyperaccumulation in infected plants (log_FC>1). Notable examples include 26-hydroxybrassinolide, campesterol, campestanol, castasterone, cathasterone, and teasterone. The final product of the pathway, brassinolide, showed substantial hyperaccumulation, with log_ fold changes of 7.47 in leaves and 7.99 in flowers following Phytoplasma infection.

Transcriptome profiling further revealed that most genes associated with brassinosteroid metabolism exhibited an upregulating trend in infected foliar tissues. These included *3-oxo-5-alpha-steroid 4-dehydrogenase 2-like* (isoform X2) and *3*β*-hydroxysteroid-dehydrogenase/decarboxylase* (isoform X2) that exhibited significant overexpression (log_FC>1). A similar trend was observed in the floral tissues, where genes like *delta(7)-sterol-C5(6)-desaturase* and *7-dehydrocholesterol reductase* exhibited significant upregulation. In contrast, *3-epi-6-deoxocathasterone 23-monooxygenase*, which catalyzes a minor branch-point step in brassinosteroid biosynthesis, was significantly downregulated in both tissues (leaves: log_FC =-0.97; flowers: log_FC =-1.90).

Interestingly, the brassinosteroid catabolic gene *cytochrome P450 734A1* (BAS1) was downregulated in both tissues, with a more pronounced suppression in floral tissues (leaves: log_FC =-0.51; flowers: log_FC =-1.02), providing a plausible explanation for the sustained accumulation of brassinosteroids during infection.

When the genes behind the brassinosteroid signalling components were examined, the receptor *Brassinosteroid Insensitive 1* (BRI1) was overexpressed in both leaf and flower tissues, with higher expression in infected flowers (log_FC = 1.476) compared to infected leaves (log_FC = 0.46). Its co-receptor, *Brassinosteroid Insensitive 1-Associated Receptor Kinase 1* (BAK1), was also upregulated in infected floral (log_FC = 1.95) and foliar (log_FC = 2.33) tissues. Additionally, the *GTP-binding protein BRASSINAZOLE INSENSITIVE PALE GREEN 2*, which is required for BR-mediated post-transcriptional regulation, was strongly enriched in both floral (log_FC = 3.95) and leaf tissues (log_FC = 1.83), further strengthening the hypothesis of tissues specific responses upon Phytoplasma infection.

## Discussion

### Phytoplasma hijacks the normal flowering machinery

Flowering in angiosperms is a highly regulated developmental process regulated by intricate intracellular signalling pathways that govern the transition of vegetative shoot apical meristem (SAM) into a floral meristem, ultimately leading to the formation of floral organs. However, in Phytoplasma-infected plants, floral tissues often exhibit virescence and phyllody, wherein floral organs are transformed into leaf-like structures. This indicates a disruption of normal floral developmental programs and suggests a phenomenon of cellular dedifferentiation followed by redifferentiation, wherein fully differentiated floral tissues revert to a meristematic state and subsequently differentiate into vegetative structures.

Previous studies have shown that this aberrant floral development is mediated by effector proteins secreted by Phytoplasmas, which are translocated through the phloem sap and induce transcriptional reprogramming in host tissues. Several such effectors have been identified, including TENGU [55, 56], SAP54 [57], SAP05, SAP11 [58], SWP1 [59], as well as Zaofeng3 [60] and Zaofeng6 [61]. Among these, SAP54, an effector of the aster yellows Phytoplasma strain AY-WB, is particularly well-characterized. It induces phyllody in *Arabidopsis thaliana* by destabilizing MADS-box transcription factors (MTFs), such as SEP2 and AP1, which are crucial for floral organ identity [57].

Present transcriptomic analysis of Phytoplasma-infected floral tissues of sesame revealed differential expression of several MADS-box transcription factors. Notably, *apetala* (AP), *agamous* (AG), and *sepallata* (SEP) family members. Especially, *AP2*-like and *SEP2*-like genes, showed significant downregulation in infected samples. These findings are consistent with earlier reports in *Catharanthus roseus* [7] and *Solanum lycopersicum* cv. Micro-Tom [62], where SEP genes were similarly repressed upon Phytoplasma infection. The suppression of *AP* and *SEP* likely contributes to the inability of infected flowers to properly develop sepals and petals, resulting in compromised floral organogenesis.

In addition to MTFs, other floral homeotic genes, such as *deficiens* (DEF) and *globosa* (GLO), which are essential for the specification of petal and stamen identity, as demonstrated in *Antirrhinum majus* <u>(Zahn *et al.*, 2005)</u>, were also significantly downregulated in infected floral tissues. These genes are orthologous to the *AP3/PI* gene family in *Arabidopsis*. Knockout mutants of *defA-1* in *A. majus* have been shown to convert petals into sepals, while *GLO* mutants exhibit sepaloid carpels. The observed repression of these genes in sesame suggests that Phytoplasma effectors interfere with the regulatory network responsible for floral organ identity, leading to profound developmental alterations and loss of floral specificity.

Interestingly, our data also revealed upregulation of several meristem identity and maintenance genes, including *clavata* (CLV), *squamosa promoter binding protein-like* (SPL), and *wuschel* (WUS) homeobox genes. This suggests that Phytoplasma infection reactivates a meristematic gene expression program in fully differentiated floral tissues. The observed antagonistic expression pattern, characterized by downregulation of floral organ identity genes and concurrent upregulation of meristem maintenance genes, may represent the molecular basis of the floral-to-vegetative retrogressive reprogramming seen in Phytoplasma-infected sesame.

Importantly, our transcriptome data did not reveal any consistent, directional changes in the expression of flowering time regulators such as *constans* (CO), *flowering locus D* (FLD), and *flowering locus k* (FLK). This suggests that while floral organ development is severely affected in infected plants, the upstream signalling pathways responsible for floral induction remain largely unperturbed. In other words, both exogenous (e.g., photoperiodic) and endogenous cues for flower initiation appear to remain functional despite the infection, allowing floral buds to form, but leading to malformed floral structures due to downstream effector-mediated reprogramming.

### Phytoplasma triggers reprogramming of Chlorophyll pathway

Although Ca. Phytoplasma–induced floral virescence is among the most prominent phenotypic symptoms observed in *S. indicum* following infection, the underlying molecular mechanisms remain poorly understood. As evident from the present study, this phenotype is characterized by the elevated accumulation of chlorophyll-a, chlorophyll-b, and biosynthetic intermediates such as protoporphyrinogen IX, coproporphyrin III, uroporphyrinogen III, 7(1)-hydroxychlorophyll-a, chlorophyllide, and 7(1)-hydroxychlorophyllide, along with a marked downregulation of chlorophyllase and pheophytinase. These molecular alterations likely explain the pronounced increase in total chlorophyll content observed in infected floral tissues. This symptom is particularly evident when the Phytoplasma titre is extremely high in flowers [7].

In contrast, the effects of Phytoplasma infection on chloroplast physiology and photosynthetic performance in leaf tissues have been extensively documented. In *Solanum lycopersicum*, infection results in severe chloroplast dysfunction, often leading to chlorophyll loss and leaf chlorosis [64].

Similarly, [18] reported that severe infection disrupts the grana and stroma lamellae, while Ahmed *et al.*, (2022), using transmission electron microscopy, showed partial degradation of both the outer and inner chloroplast membranes. Phytoplasma infection is also known to impair PSII efficiency and photosynthetic performance in leaves, as reported in *Pennisetum purpureum* [65], *Malus pumila* [66], and *Prunus avium* [21]. In our present study, transcriptome analysis of infected leaf tissues revealed upregulation of chlorophyll biosynthetic genes, which could possibly be a compensatory response. As, it was accompanied by a significant increase in chlorophyll catabolism, indicated by elevated expression of chlorophyllase and pheophytinase and enhanced accumulation of chlorophyll breakdown metabolites including pheophorbide-a, primary fluorescent chlorophyll catabolite, and red fluorescent catabolites etc. These findings suggest a dynamic equilibrium between chlorophyll synthesis and degradation in infected leaf tissues, resulting in a relatively stable overall chlorophyll level in leaf tissues, in contrast to the pronounced increase seen in floral tissues. The possible reason is, unlike previous studies reporting a substantial decline in chlorophyll a and b in infected leaves [66–68] our spectrophotometric data did not show a marked reduction in total chlorophyll content; and it could be due to differences in chlorophyll turnover dynamics.

### Oxidative burst serves in triggering defence mechanisms

Reactive oxygen species (ROS) play a dual role in plants, functioning both as signalling molecules under normal physiological conditions and as mediators of stress [69]. The ROS are scavenged by both enzymatic and non-enzymatic antioxidant defense systems. The enzymatic system comprises superoxide dismutase (SOD), catalase (CAT), and peroxidases (POXs), while the non-enzymatic system includes antioxidants such as glutathione, ascorbic acid, carotenoids, and flavonoids [70, 71]. These antioxidants neutralize free radicals, and assays such as DPPH, ABTS, FRAP, and superoxide (SO) scavenging provide an estimate of the antioxidant capacity of a tissue [72]. In the present study, antioxidant content increased in both foliar and floral tissues upon Ca. Phytoplasma infection, as evidenced by TAA, DPPH, ABTS, FRAP, and SO assays, suggesting an enhanced oxidative response to infection.

At the transcriptome level, the most notable change was the strong upregulation of SOD genes in both floral and leaf tissues following infection. SOD catalyzes the dismutation of superoxide anions into H_O_, and this increase was consistent with spectrophotometric SO scavenging data. Similar increases in SOD activity have been reported in *Citrus aurantifolia* [73], *Citrus sinensis* [68], *Chrysanthemum coronarium* [74], and *Vigna radiata* [67].

Peroxidases, which function downstream of SOD by detoxifying H_O_, were also markedly upregulated in infected leaf and flower tissues. It included ascorbate peroxidase (APOX) and cationic peroxidases, as confirmed by both transcriptomic and spectrophotometric data (GPOX and APOX activity). Previous studies have similarly reported increased peroxidase activity during Phytoplasma infection [65, 67, 68, 75, 76]. Infected floral tissues showed enhanced expression of APOX, lignin-forming anionic peroxidases, and multiple cationic peroxidases, while leaf tissues displayed upregulation of multiple APOX and cationic peroxidase isoforms. A few peroxidase transcripts, however, were downregulated in infected floral tissues.

APOX enzymes detoxify H_O_ using ascorbic acid as a reducing substrate. Their upregulation is consistent with earlier reports in *Catharanthus roseus* [75] and *Citrus sinensis* [68]. Asudi *et al.*, (2021)also reported increased ascorbic acid content in *Pennisetum purpureum* during Napier grass stunt Phytoplasma infection, supporting the observed enhancement of APOX activity. Cationic peroxidases, including guaiacol peroxidase, were also universally upregulated across all tissues in our study. These enzymes can polymerize phenolic substrates, promoting lignification. Similarly, lignin-forming anionic peroxidases, upregulated in infected floral tissues, catalyze the polymerization of phenolics during H_O_ detoxification, leading to lignin deposition [77–79]. This is corroborated by our histochemical phloroglucinol-HCl staining, which revealed higher lignin deposition in infected shoots, and by the overaccumulation of syringin, a syringyl lignin precursor, in infected flowers [12].

Many of these anionic peroxidases are also involved in cell wall stiffening via suberization in addition to lignification [77–81]. Suberin is a complex biopolymer composed of a phenolic domain bound to the cell wall and aliphatic components covalently linked to this phenolic domain [82]. Its biosynthesis involves dehydrogenative polymerization of hydroxycinnamic acids, catalyzed by anionic peroxidases [83]. In our metabolomic study, 4-hydroxycinnamic acid accumulated significantly in floral tissues (log_FC = 0.98), and was also elevated in leaves. The concurrent upregulation of lignin-forming anionic peroxidases and accumulation of lignin/suberin precursors suggest that enhanced lignification and suberization could act as structural defenses, restricting Phytoplasma movement from source (infection reservoirs) to sink tissues.

In contrast to peroxidases, catalase (CAT) gene expression remained largely unchanged in floral tissues and was only marginally upregulated in infected leaves. Since CAT is mainly localized in peroxisomes and chloroplasts [84, 85], and Phytoplasma infection is known to disrupt organellar integrity [86, 87], its activity may be inherently limited under such conditions. It aligns with previous studies reporting reduced CAT activity in *Vigna radiata* [67] and *Catharanthus roseus* [75] during Phytoplasma infection.

An interesting tissue-specific trend was observed for polyphenol oxidase (PPO). The activity and gene expression of PPO were higher in the infected floral tissues but negligible in leaves. Earlier studies reported reduced PPO activity in leaves during Phytoplasma infection (*Vigna radiata*: Hameed *et al.*, (2017); *Catharanthus roseus*: [75]. The PPOs are predominantly localized in chloroplasts, either in the lumen or associated with thylakoid membranes near Photosystem I and II [88–92]. The observed decline in leaf PPO activity can thus be attributed to chloroplast degradation, a hallmark of Phytoplasma infection [86, 87]. Conversely, Phytoplasma-induced floral virescence promotes chlorophyll accumulation and the development of organized chloroplasts in floral tissues, which was confirmed by our spectrophotometric data. It likely explains the elevated PPO activity observed in infected flowers.

### Octadecanoid pathway alteration upon Phytoplasma infection

Activation of the octadecanoid pathway is among the earliest defense responses in plant tissues exposed to biotic stress. This response is typically triggered by a rapid oxidative burst of reactive oxygen species (ROS) following pathogen attack, which induces the release of 9,12,15-octadecatrienoic acid (α-linolenic acid) from membrane-associated polar lipids through enzymatic or non-enzymatic mechanisms. Depending on the nature of the external biotic stressors, the pathway ultimately leads to the biosynthesis of jasmonic acid (JA) [84], green leaf volatile oxylipins [93–95], or so-called “death acids” [96, 97].

Previous studies have shown that JA can activate the phenylpropanoid pathway by inducing phenylalanine ammonia-lyase (PAL) activity, as demonstrated in *Hypericum perforatum* L. [98] and *Oryza sativa* [99]. Similarly, methyl jasmonate (MeJA)-mediated activation of the phenylpropanoid pathway has been reported in *Actinidia chinensis* [100]. While JA and MeJA signalling are well established in plant defense against necrotrophic pathogens and herbivorous insects [101–104], their effectiveness against biotrophic pathogens such as Phytoplasmas remains less defined [105].

Conflicting reports exist regarding JA dynamics in Phytoplasma infections. [25] observed significant downregulation of JA and its bioactive conjugate JA-Ile in the apical meristems of *S. indicum* infected with *Ca.* Phytoplasma onion yellows strain. In contrast, <u>Karan *et al.*, (2025.)</u>documented overexpression of the jasmonate-O-methyltransferase gene in *S. indicum* leaves upon *Ca.* Phytoplasma asteris infection. Similarly, <u>Mardi *et al.*, (2015)</u> reported overexpression of the same gene in lima bean leaves infected with *Ca.* Phytoplasma aurantifolia. Notably, both Verma et al. (2025) and Mardi et al. (2015) reported downregulation of early octadecanoid pathway genes, including lipoxygenase (*LOX*), allene oxide synthase (*AOS*), and allene oxide cyclase (*AOC*) family members.

In the present study, we observed marginally higher JA accumulation in both leaf (log_2_FC: 0.28) and floral tissues (log_2_FC: 0.51) upon infection, with a comparatively greater increase in floral tissues. Methyl jasmonate levels were somewhat higher in infected leaves (log_2_FC: 0.51) but significantly depleted in infected floral tissues (log_2_FC:-1. 17), concerning the respective controls. Transcriptomic data revealed downregulation of key JA biosynthetic genes (e.g., AOS and AOC family) in leaf tissues. Given that these early biosynthetic steps occur in the chloroplast, their reduced activity may be linked to compromised chloroplast function. Indeed, reduced PPO activity in previous studies suggested impaired chloroplast performance, and <u>Wei *et al.*, (2022)</u>reported chloroplast dysfunction and structural degradation following Phytoplasma infection. Such dysfunction likely explains the reduced activity of early octadecanoid pathway genes and the marginal JA accumulation observed in infected leaves. Conversely, MeJA biosynthesis occurs in the cytosol, which may account for its relatively higher accumulation in leaf tissues.

In the floral tissues, early pathway genes such as *AOS* and *12-OPDA* reductase were upregulated, likely reflecting the functional status of chloroplasts in flowers, possibly linked to floral virescence symptoms. However, the peroxisomal “late” pathway genes were downregulated, potentially limiting JA accumulation despite active early steps. The relatively higher JA levels in flowers may also contribute to a stronger impact on the phenylpropanoid pathway in floral tissues compared to leaves.

The observed tissue-specific differences in JA accumulation can be attributed to multiple factors: Firstly, organelle functionality, i.e., variations in chloroplast, peroxisome, and cytosolic function between tissue types; secondly, precursor availability, i.e., Infected floral tissues exhibited greater α-linolenic acid accumulation than leaves, providing a larger substrate pool for JA biosynthesis. Finally, branch pathway flux, i.e., infected leaves showed substantial accumulation of green leaf volatiles (GLVs) such as 3-hexenyl acetate and 3-hexen-1-ol, along with their biosynthetic intermediates [13(S)-HPOT and 9S-HpOTrE], which are terminal products of an alternative branch of the octadecanoid pathway. In contrast, these metabolites were significantly downregulated in infected floral tissues, potentially contributing to higher JA retention in flowers.

The GLVs are recognized as a key frontline defense against a range of biotrophic and necrotrophic pathogens and herbivorous insects <u>Matsui and Engelberth, (2022)</u>. While not previously reported in Phytoplasma patho-systems, López-Gresa et al. (2018) showed that antisense suppression of 3-hexenyl acetate in tomato increased susceptibility to *Pseudomonas syringae*. In the context of Phytoplasma infection, <u>Ferreira *et al.*, (2023)</u> first reported GLV overaccumulation in cabbage palms (*Sabal palmetto*), infected with lethal bronzing Phytoplasma. The defensive role of GLVs against Phytoplasmas may be indirect but ecologically significant, as Phytoplasma diseases are insect vector-borne. Numerous studies on planthoppers and other sap-sucking insect vectors have documented strong induction of GLVs following herbivory [110–112], and these compounds act as potent repellents against sap-sucking insects such as the brown planthopper [110, 112, 113].

Taken together, these findings suggest that *S. indicum* mounts a tissue-specific, multi-layered defense strategy upon Phytoplasma infection, in which variations in JA/MeJA dynamics, GLV production, and organelle function collectively shape the outcome of the plant–pathogen interaction.

### Phytoplasma alters the phenylpropanoid pathway

Phenylpropanoids play a central role in strengthening plant defense mechanisms against biotic stressors. They generate diverse metabolites, including phenolics, which function as substrates for stress-responsive peroxidases (as discussed earlier) or undergo enzymatic polymerization to form lignin. Lignin deposition enhances cell wall rigidity, thereby establishing a mechanical barrier to pathogen invasion. A specific subset of phenylpropanoids, the phenolic acids, particularly hydroxycinnamic acids, contribute to cell wall suberization, another defense strategy that reinforces barriers against pathogen spread.

The present study demonstrates that Phytoplasma infection leads to the accumulation of multiple phenylpropanoid pathway metabolites and the overexpression of genes associated with this pathway in both examined tissues. Our previous work had already reported the upregulation of the phenylpropanoid pathway as a significant consequence of Phytoplasma infection in sesame leaves (Banerjee and Gangopadhyay 2023). The current tissue-specific analysis further reveals that floral tissues exhibit a much stronger alteration in phenylpropanoid metabolism compared to leaves. This observation is consistent with prior reports documenting that Phytoplasma titre is generally higher in floral tissues [114], likely triggering stronger stress responses and consequently a greater impact on the phenylpropanoid pathway.

Similar responses have been observed in other plant species. For instance, Phytoplasma infection of *Camptotheca acuminata* showed upregulation of both differentially expressed genes (DEGs) and metabolites of the phenylpropanoid pathway (Qiao et al., 2023). Comparable upregulation has also been documented in *Citrus aurantifolia* (Mardi et al., 2015), *Paulownia fortunei* (Fan et al., 2015), *Malus domestica* (Patui et al., 2013), and *Catharanthus roseus* (Choi et al., 2004). However, more recent work on S. indicum reported downregulation of the phenylpropanoid pathway in response to infection with Phytoplasma strains such as onion yellows [25] and asteris <u>(Karan *et al.*, 2025</u>). These contrasting results suggest that different Phytoplasma strains (present findings with strain aurantifolia) may elicit distinct host responses.

The primary contribution of the phenylpropanoid pathway to pathogen resistance is through lignification, a process that modifies the cell wall. In the present study, increased levels of p-coumaric acid, caffeoyl alcohol, cis-caffeic acid, sinapic acid, sinapyl alcohol, and syringin, alongside upregulation of rate-limiting genes such as PAL and 4CL, and peroxidase genes involved in polymerization, indicate enhanced lignin biosynthesis, particularly syringyl lignin. Similarly, it was reported that PEROXIDASE 51, which catalyzes lignin monomer polymerization, was upregulated in Phytoplasma-infected jujube (*Ziziphus jujuba*), and its overexpression in *Arabidopsis* significantly improved Phytoplasma resistance. Lignin fortifies defense against insect-borne bacterial pathogens in multiple ways: (i) it resists microbial degradation [115] restricting pathogen spread through lignification of infected tissues; and (ii) it reduces palatability for sap-sucking insect vectors by hardening plant tissues. Supporting this, <u>He *et al.*, (2020)</u> demonstrated that overexpression of OsPALs in rice enhanced lignin biosynthesis and reduced susceptibility to brown planthopper (BPH), whereas gene silencing had the opposite effect. Similarly, an OsSLR1-silenced rice line, characterized by elevated phenolic acid and lignin levels, showed improved resistance to BPH [117].

In addition to lignification, our results suggest that suberization may also be involved in the defense response to Phytoplasma infection. KEGG pathway enrichment analysis, together with the accumulation of hydroxycinnamic acids and peroxidase overexpression, indicates that suberization is particularly prominent in leaf tissues. Deposition of ligno-suberin in vascular tissues is a well-established defense mechanism against pathogens that colonize vascular tissues [118]. They further showed that induced ligno-suberin vascular coating and hydroxycinnamic derivatives restricted *Ralstonia solanacearum* colonization in tomato vasculature [119]. Likewise, [120]) reported suberization of vascular tissues in Ulmus minor as a key defense against *Ophiostoma novo-ulmi*.

Among other tissue-specific changes, the overaccumulation of lignans such as pinoresinol and secoisolariciresinol in infected leaves was notable, consistent with previous findings from our group (Banerjee and Gangopadhyay, 2024). Lignans are multifunctional defense compounds with demonstrated antifeedant activity [121], insect growth inhibitory effects [122], and endocrine-disrupting properties [123] against hymenopteran, dipteran, and hemipteran insects. Their higher accumulation in infected leaves compared to flowers may reflect an adaptive response, since BPH primarily targets leaf tissues for phloem sap extraction, with no reports of BPH feeding on flowers. We therefore hypothesize that increased pinoresinol accumulation represents a defensive strategy to reduce BPH infestation and consequently limit Phytoplasma spread.

### Phytoplasma trigger brassinosteroid biosynthesis pathway

One of the most consistent changes observed in both tissues following infection was the upregulation of the brassinosteroid (BR) biosynthesis pathway, at least at the metabolic level. Transcriptome data also supported this metabolic trend to some extent. The impact of *Ca.* Phytoplasma *aurantifolia* infection on the BR biosynthesis pathway has been previously documented by <u>Mardi *et al.*, (2015)</u> in *Citrus aurantifolia* L., who reported that a wide array of unigenes associated with BR biosynthesis were upregulated at the transcriptomic level upon infection. Similar findings were later described by <u>Wang *et al.*, (2018)</u> in Ziziphus jujuba infected with *Ca*. Phytoplasma *ziziphin*. We found both *BRI1* and *BAK1* to be upregulated upon infection and both Mardi et al. (2015) in *C. aurantifolia* and <u>Yan *et*</u> *<u>al.</u>*<u>, (2019)</u> in *Paulownia fortunei* reported a significant upregulation of the BRI1-Associated Receptor Kinase 1 (BAK1) gene upon Phytoplasma infection, which they hypothesized that *BAK1* may act as a key mediator enabling host plants to withstand Phytoplasma invasion.

Brassinosteroids are primarily recognized as plant growth regulators involved in diverse developmental processes; however, their role in biotic stress responses is increasingly acknowledged [126, 127]. Growing evidence suggests that BRs function as important mediators of defense crosstalk during plant–pathogen interactions, coordinating with hormones such as gibberellic acid (growth-related), salicylic acid (defense-related), and abscisic acid in the context of PAMP-triggered immunity (growth-defense continuum). BR signalling has also been reported to fine-tune cellular redox balance and secondary metabolism [128].

For instance, <u>Nakashita *et al.*, (2003)</u> demonstrated that *Nicotiana tabacum* cv. *Xanthi* infected with TMV exhibited a marked increase in BR levels, implicating their role in plant defense. In subsequent studies, the exogenous application of brassinolide enhanced disease resistance against TMV as well as bacterial pathogens such as *Magnaporthe grisea* and *Xanthomonas oryzae* pv. *oryzae*. This phenomenon was termed steroid hormone-mediated disease resistance (BDR).

Although the precise role of brassinosteroids in Phytoplasma infections remains poorly understood, Mardi et al. (2015) hypothesized that BR-related defense mechanisms may involve the induction of PR1 expression and activation of salicylic acid biosynthesis, thereby contributing to pathogen resistance.

## Conclusion

In conclusion, our multi-omics investigation provides a comprehensive understanding of how sesame plants respond to *Ca*. Phytoplasma *aurantifolia* infection, with a particular focus on exploring tissue-specific responses in leaf and floral tissues (Fig. 10). Phytoplasma infection induces retrogressive morphogenesis, converting floral tissues into leafy structures, which is associated with reduced expression of floral organ identity genes and concomitant upregulation of meristem-maintaining genes.

**Fig. 10:**
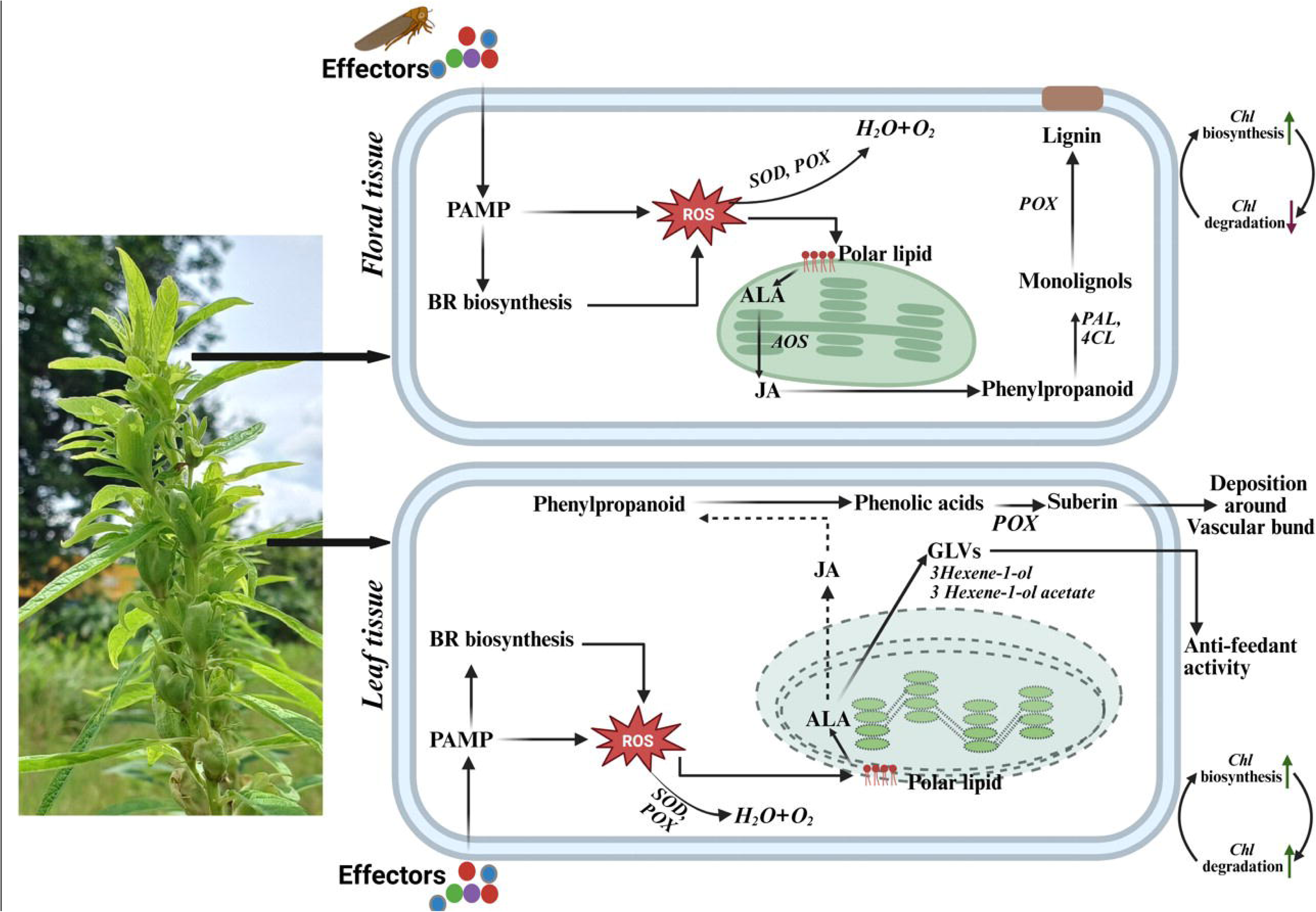
Graphical illustration of the proposed mechanism underlying tissue-specific defense responses of sesame plants against *Ca.* Phytoplasma infection.

We demonstrate that Phytoplasma infection triggers profound tissue-specific reprogramming at both metabolic and transcriptional levels. Floral tissues, in particular, exhibited stronger alterations than leaf tissues, consistent with their higher pathogen titre and phenotypic susceptibility. Essential pathways, including chlorophyll metabolism, phenylpropanoid biosynthesis, brassinosteroid biosynthesis, octadecanoid metabolism, redox homeostasis, and cutin, suberin wax biosynthesis were significantly influenced. These molecular changes were validated by qRT-PCR, spectrophotometry, histochemical assays, radical scavenging assays, and antioxidant enzyme activity measurements, providing multiple layers of evidence. In comparison with the previous studies, we hypothesize that the alterations induced by *Ca*. Phytoplasma *aurantifolia* in sesame may differ from those caused by other Phytoplasma strains in same host plants, highlighting the strain-specific nature of host–pathogen interactions. Therefore, detailed investigations are essential to fully elucidate the multidimensional networks underpinning Phytoplasma pathogenesis. Overall, our study provides an integrative picture of the metabolic and transcriptional reprogramming in floral and leaf tissues of sesame, thereby laying a foundation for a better understanding of Phytoplasma infections and the development of potential management strategies.

## Author Contribution

SB performed the wet lab experiments, analysed the data, and wrote the manuscript. GG conceived the study, designed, supervised, and contributed to the manuscript writing.

## Acknowledgement

We acknowledge Mr Santu Pal, Indian Institute of Chemical Biology, for assistance in LC-MS/MS. We also acknowledge the technical assistance of Mr Jadab Ghosh, Mrs Kaberi Ghosh, and Mr Swarnava Das, Department of Biological Sciences, BI, in a few laboratory experiments.

## Clinical trial number

Not applicable.

## Data Availability Statement

The Illumina sequencing data sets generated are available in Sequence Read Archive, NCBI under the accession number of PRJNA1307559 (https://www.ncbi.nlm.nih.gov/sra/PRJNA1307559).

## Ethics approval and consent to participate

Not applicable

## Consent for publication

Not applicable.

## Abbreviation

13(S)-HPOT: (9Z,11E,13S,15Z)-13-hydroperoxyoctadeca-9,11,15-trienoic acid
4CL2: 4-coumarate--CoA ligase 2
7-HCideAR: 7(1)-Hydroxychlorophyllide a
9,10-EOT: 9,10-Epoxyoctadecatrienoic acid
AG: Agamous
ANR: Anthocyanidin Reductase
AOC: Allene oxide cyclase
AOS1: Allene oxide synthase
AP: Apetala
APX: L-ascorbate peroxidase
C4H: Trans-Cinnamate 4-Monooxygenase
CAD: Cinnamyl-alcohol dehydrogenase
CAO: Chlorophyllide a oxygenase
CAT: Catalase
CatPER: Cationic peroxidase
CBR: Chlorophyll(ide) B reductase
CCoAOMT: Caffeoyl-CoA O-methyltransferase
CCR1: Cinnamoyl-CoA reductase 1
CCR2: Cinnamoyl-CoA reductase 2
CHCI: Magnesium-chelatase subunit chlI
Chl: Chlorophyll
CHLD: Magnesium-chelatase subunit CHLD
CHLG: Chlorophyll synthase
CHLH: magnesium-chelatase subunit CHLH
Chlide: Chlorophyllide
CLH2: Chlorophyllase-2
CLV: Clavata
COL: Constans-like
COMT: Caffeic acid 3-O-methyltransferase
COPRO III: Coproporphyrin III
Coprogen III: Coproporphyrinogen III
CPOX: Coproporphyrinogen-III oxidase 1
DEF1: Deficiens1
ECH2: Enoyl-CoA hydratase 2
FAD4: Fatty acid desaturase 4
FLD: Flowering locus D
GLO: Globosa
GLO: Lactoylglutathione lyase
GPX: Glutathione peroxidase 2
GST: Glutathione S-transferase
GSTU17-like: Glutathione S-transferase U17-like
HAGH: Hydroxyacylglutathione hydrolase
HCAR: 7(1)-Hydroxychlorophyll a
HpOTrE: 12-hydroperoxy-9Z,13E,15-octadecatrienoic acid
KAT2: 3-ketoacyl-CoA thiolase 2
LDOX: Leucoanthocyanidin Dioxygenase
Lip: Lignin-forming anionic peroxidase
LOX2.1: Linoleate 13S-lipoxygenase 2-1
LOX3.1: Linoleate 13S-lipoxygenase 3-1
LOX5: Linoleate 9S-lipoxygenase 5
LOX6: Lipoxygenase 6
Mg-Proto: Magnesium protoporphyrin
MGST3: Microsomal glutathione S-transferase 3
MPEC: Mg-protoporphyrin IX monomethyl ester cyclase
NDC: NAD(P)H-ubiquinone oxidoreductase C1
NDUFA: NAD(P)H-ubiquinone oxidoreductase A1
NDUFAF: NADH dehydrogenase (ubiquinone) complex I, assembly factor 6
NDUFB: NADH dehydrogenase [ubiquinone] 1 beta subcomplex subunit B-like
NDUFB: NAD(P)H-ubiquinone oxidoreductase B
NDUFS: NADH dehydrogenase [ubiquinone] iron-sulfur protein
NDUFV: NADH dehydrogenase [ubiquinone] flavoprotein
NQO: NAD(P)H:quinone oxidoreductase-like
OPR11: 12-oxophytodienoate reductase 11
PAL: Phenylalanine ammonia-lyase
PBG: Porphobilinogen
PBGD: Porphobilinogen deaminase
Pchlide: Protochlorophyllide
PER: Peroxidase
PhA: Pheophorbide a
PHGPX: Phospholipid hydroperoxide glutathione peroxidase 1
PORA: Protochlorophyllide reductase
PPG: Protoporphyrinogen IX
PPH: Pheophytinase
PPOX: Protoporphyrinogen oxidase
PPOX: Polyphenol oxidase
SEP: Sepallata
SOD: Superoxide dismutase
SPL: Squamosa promoter-binding-like protein
UROD: Uroporphyrinogen decarboxylase
Urogen III: Uroporphyrinogen III

