## Supplementary material for "*Candidatus* Phytoplasma-induced Retrogressive Morphogenesis in Sesame (*Sesamum indicum* L.): Tissue-Specific Metabolic and Transcriptomic Reprogramming": Table S1-S2

Table S1: Table for Nested PCR

| **Name** | **Primer sequence** | **Annealing temperature** |
| --- | --- | --- |
| P1 | AAGAGTTTGATCCTGGCTCAGGATT | 55°C |
| P7 | CGTCCTTCATCGGCTCTT | 55°C |
| R16F2n | GAAACGACTGCTAAGACTGG | 60°C |
| R16R2 | TGACGGGCGGTGTGTACAAACCCCG | 60°C |

Table S2: Primers for real time PCR

| **Sl. No** | **Gene name** | **Primer sequence (5′→3′)** | **Amplicon size (bp)** | **Accession number** |
| --- | --- | --- | --- | --- |
| 1 | Sepalata-like 1 | F- ACAAGGACGCAGGTTATGCT  R- ACCCATGATTGCTGAAGCTGA | 134 | XM_011103957.2 |
| 2 | Apetala 1 | F- GAAGCACACCAATCTCCCCAT  R- TCCTTTATGCCCAACTGTCCC | 134 | XM_020691960.1 |
| 3 | Flowering locus D | F- GTACCGAAGTCGAAAGCAGC  R- CGTTGCTGCCGACTTTGCTT | 121 | XM_011096042.2 |
| 4 | WUSCHEL-related homeobox 1 | F- GTTGTGGTTAGTTCGCGCTG  R-TGTGCGGTAATGTGCTGGAT | 116 | XM_011085944.2 |
| 5 | Phenylalanine ammonia lyase | F- AAAGAAGGGCTTGCACTGGT  R- CGGTGAACTCAGGCTTTCCA | 148 | XM_011079036.2 |
| 6 | Allene oxide cyclase | F- ACGAGATGAACGAGCGAGAC  R- TATGTCAGATACGCGCCCTG | 256 | XM_011080147.2 |
| 7 | Allene oxide synthase 1 | F-CCAACCACGCCAAACTCAAG  R- GGCCAAGAAATTGAAGGCGG | 182 | XM_011076172.2 |
| 8 | 4-coumarate--coa ligase 1 | F- TGCAAAACTTGGACAGGGGT  R- ATTTCTCCGGGCTGGTTACG | 186 | XM_011074951.2 |
| 9 | Polyphenol oxidase I | F- GACCAGAAACCCTGACCCAC  R- TGGCAGTAGAACAACGGGTC | 261 | XM_011084554.2 |
| 10 | Linoleate 13S-lipoxygenase 2-1 | F- TGTCTACAACGATCTCGGCG  R- AAATGCCTCGTCTCTTGGCA | 175 | XM_011081940.2 |
| 11 | Cinnamyl alcohol dehydrogenase 1 | F- GGGGGATTTGCTGGTGCTAT  R- CGAGACCCAAAATACCGCCT | 172 | XM_011089904.2 |
| 12 | Anthocyanidin reductase | F- TGGGGCTTCTACAGGGTCTT  R- CTTAACCCCACCCATAGCCG | 203 | XM_011095240.2 |
| 13 | Leucoanthocyanidin dioxygenase | F- ACTGCACTTGCCTGATAAACC R- TGCCGAAAGCATGACTCCAT | 171 | XM_011094135.2 |
| 14 | Lignin-forming anionic peroxidase | F- AAGGGACTCTACCACAGCCA  R- TATGTGCTCCAGAAAGGGCG | 140 | XM_011072295.2 |
| 15 | Cinnamoyl-CoA reductase 1 | F- GCCAGAAATGTGATACGGGC  R- GTCCACCCCAAACTCCTCAG | 228 | XM_011075026.2 |
| 17 | Actin | F-ATGGAAGCTGCAGGCATTCA  R-TGATGCTAGGATTCACCTCC | 250 | XM_011079162.2 |
