## Supplementary material for "*Candidatus* Phytoplasma-induced Retrogressive Morphogenesis in Sesame (*Sesamum indicum* L.): Tissue-Specific Metabolic and Transcriptomic Reprogramming": Table S4

| Sample | Nano reading  (ng/µl) | RIN value | Total Number of reads | Total number of bases | No. of CDS | Mean CDS length (bp) | Total number of annotated CDS |
| --- | --- | --- | --- | --- | --- | --- | --- |
| CF1 | 228.0 | 6.7 | 18,732,964 | 5,657,355,128 | 33,397 | 1,159 | 26,548 |
| CF2 | 224.0 | 6.6 | 15,937,967 | 4,183,266,034 | 28,882 | 1,116 | 22,903 |
| CF3 | 228.7 | 6.7 | 14,697,808 | 4,438,738,016 | 28,061 | 1,138 | 22,323 |
| CL1 | 1423.7 | 5.7 | 12,725,428 | 3,843,079,256 | 26,544 | 1,109 | 21,081 |
| CL2 | 901.6 | 5.5 | 15,717,183 | 4,746,589,266 | 30,834 | 1,160 | 24,477 |
| CL3 | 328.6 | 5.7 | 15,215,529 | 5,595,089,758 | 30,008 | 1,158 | 23,873 |
| IF1 | 575.0 | 6.6 | 13,447,691 | 4,061,202,682 | 29,912 | 1,149 | 23,838 |
| IF2 | 585.9 | 6.7 | 16,079,679 | 4,856,063,058 | 31,429 | 1,158 | 24,992 |
| IF3 | 526.9 | 6.7 | 15,190,792 | 4,587,619,184 | 30,295 | 1,149 | 24,043 |
| IL1 | 250.8 | 5.6 | 14,217,596 | 4,293,713,992 | 27,036 | 1,141 | 21,469 |
| IL2 | 336.9 | 5.5 | 14,437,164 | 4,360,023,528 | 28,065 | 1,146 | 22,326 |
| IL3 | 328.6 | 5.4 | 18,366,871 | 5,546,795,042 | 31,206 | 1,146 | 24,595 |

Table S4: RNA quality, quantification and read statistics of the sample data

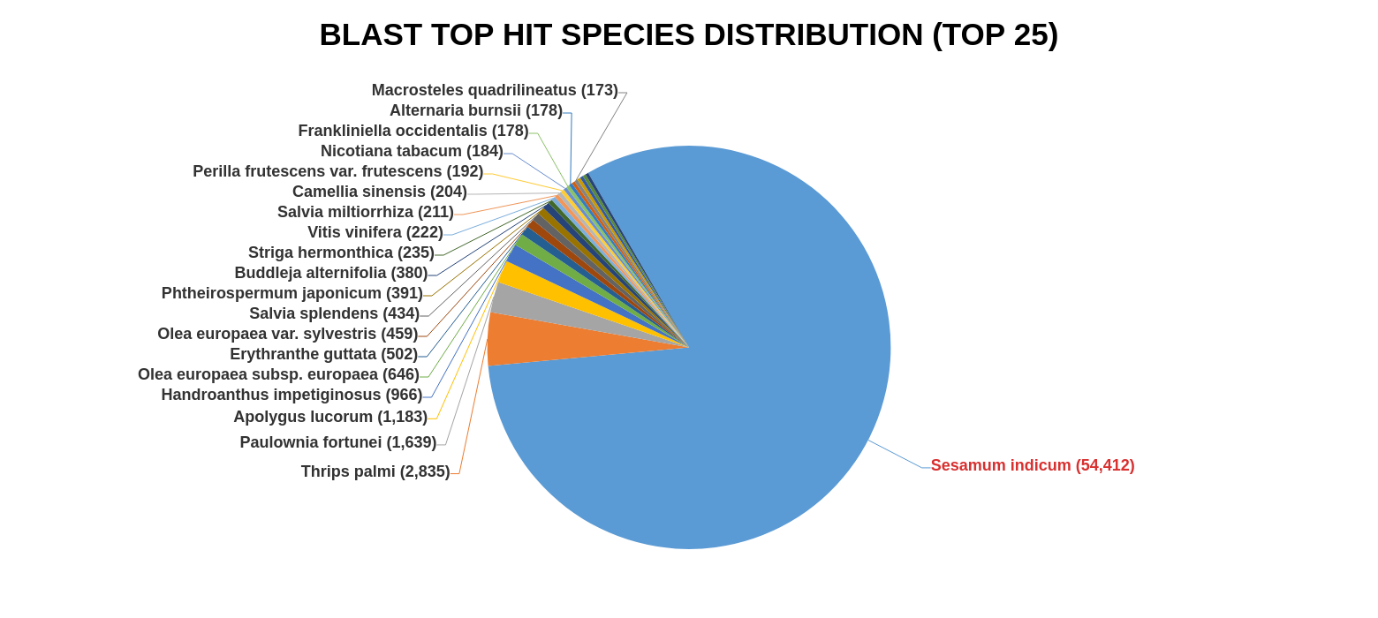

Figure S1: Top Blast Hit species distribution of pooled CDS
