## Supplementary material for "*Candidatus* Phytoplasma-induced Retrogressive Morphogenesis in Sesame (*Sesamum indicum* L.): Tissue-Specific Metabolic and Transcriptomic Reprogramming": Table S6

|  | Foliar tissues | | | Floral tissues | | |
| --- | --- | --- | --- | --- | --- | --- |
|  | CL | IL | p-value | CF | IF | p-value |
|  | Plant secondary metabolites | | | | | |
| Total Phenolic Content | 60.5381±6.739 | 84.2180±6.7365 | p=0.000536 | 19.3193±2.018 | 58.7939±14.558 | p=0.00032 |
| Total Flavonoid Content | 51.8276±6.838 | 65.3656±6.652 | p=0.00625 | 15.2868±2.011 | 43.7021±4.186 | p=1.867E-07 |
| Total Tannin Content | 8.07234±0.768 | 14.2476±2.532 | p =0.000805 | 3.57691±1.372 | 6.64285±1.174 | p=0.0052 |
|  | Plant physiological parameters | | | | | |
| Total chlorophyll content | 50.81±4.673 | 56.75±3.822 | p=0.129 | 5.38±1.417 | 49.15±4.673 | p=4.004E-08 |
| Total carotenoid content | 1.910±0.999 | 1.999±1.315 | p =0.907 | 0.242±0.102 | 3.021±0.849 | p =8.94E-05 |
|  | Activity of stress enzymes | | | | | |
| PPO | 1.350±1.1749 | 0±0 | NA | 3.470±2.578 | 6.051±1.552 | p=0.188 |
| APOX | 12.968±2.529 | 13.096 ±1.615 | p=0.9445 | 12.869±2.947 | 29.388±1.924 | p=0.0012 |
| GPOX | 0.2234±0.0418 | 24.636±4.178 | p=6.77E-05 | 5.272±0.228 | 17.937±2.635 | p=0.00115 |
|  | Antioxidant content | | | | | |
| TAA | 10.9001±1.882 | 16.6122±1.022 | p=0.0003 | 3.7313±1.815 | 9.6942±2.373 | p=0.0021 |
| DDPH | 11.6993±2.325 | 16.3954±1.776 | p=0.0071 | 6.6991±1.437 | 11.093±1.851 | p= 0.0030 |
| ABTS | 9.7128±1.605 | 12.8996±1.212 | p=0.0075 | 3.3427±1.551 | 7.3595±1.602 | p =0.0038 |
| FRAP | 18.575±1.879 | 28.5166±3.232 | p=0.0003 | 4.2060±1.373 | 14.856±2.641 | p=4.37E-05 |
| SO | 18.0177±2.417 | 21.3247±2.475 | p=0.065 | 8.0646±2.417 | 15.479±2.348 | p=0.00116 |

Table S6: Spectrophotometric data pf plant secondary metabolites, plant physiological parameters, activity of stress enzymes and antioxidant content
